# Enhancer identification and activity evaluation in the red flour beetle, *Tribolium castaneum*

**DOI:** 10.1101/199729

**Authors:** Yi-Ting Lai, Kevin D. Deem, Ferran Borràs-Castells, Nagraj Sambrani, Heike Rudolf, Kushal Suryamohan, Ezzat El-Sherif, Marc S. Halfon, Daniel J. McKay, Yoshinori Tomoyasu

## Abstract

Evolution of *cis*-properties (such as enhancers) often plays an important role in the production of diverse morphology. However, a mechanistic understanding is often limited by the absence of methods to study enhancers in species outside of established model systems. Here, we sought to establish methods to identify and test enhancer activity in the red flour beetle, *Tribolium castaneum*. To identify possible enhancer regions, we first obtained genome-wide chromatin profiles from various tissues and stages of *Tribolium* via FAIRE (Formaldehyde Assisted Isolation of Regulatory Elements)-sequencing. Comparison of these profiles revealed a distinct set of open chromatin regions in each tissue and stage. Second, we established the first reporter assay system that works in both *Drosophila* and *Tribolium*, using *nubbin* in the wing and *hunchback* in the embryo as case studies. Together, these advances will be useful to study the evolution of *cis*-language and morphological diversity in *Tribolium* and other insects.

## INTRODUCTION

Insects display some of the greatest diversity of morphology found amongst eukaryotic taxa, offering a variety of opportunities to investigate molecular and developmental mechanisms underlying morphological evolution. Decades of studies in evolutionary developmental biology (evo-devo) have revealed that changes in gene regulatory networks (GRNs) have been a major driving force in the production of the diverse morphology seen in insects as well as in other taxa (Carroll, 2008; Carroll et al., 2005). In general, a GRN can be divided into two components: *trans* and *cis*. *trans* components are transcription factors (TFs) and their upstream regulators (such as signal transduction pathways) that provide instructive cues for patterning and differentiation to the tissues where they are expressed. In contrast, *cis* components are non-coding DNA elements (*i.e. cis*-regulatory elements, CREs) that gather and process the upstream *trans* information and determine the spatial and temporal expression of the genes downstream in the genetic pathway. Changes in both *cis* and *trans* components have been implicated in morphological evolution (Carroll, 2008; Carroll et al., 2005; Halfon, 2017).

By embracing unparalleled genetic tools, both *cis* and *trans* factors have been analyzed in great detail in the fruit fly, *Drosophila melanogaster*. The accumulated knowledge obtained from *Drosophila* studies can be used as a reference (i.e. the *Drosophila* paradigm) when studying other insects and identifying the changes in GRNs responsible for morphological evolution. RNA interference (RNAi)-based gene knock-down techniques have allowed for an investigation of the *trans* properties involved in development and their evolutionary conservation/diversification in various insects (Belles, 2010). However, the lack of a reliable method to identify *cis* properties in non-*Drosophila* insects has made it difficult to study the evolution of *cis* properties beyond *Drosophila* species, even though it is equally important to study *cis* properties to gain a more comprehensive view of changes in GRNs that contributed to morphological evolution.

The major difficulty in identifying CREs, such as enhancers, stems from the labile nature of *cis* properties. The genes that code for *trans* factors important for development are usually evolutionarily well-conserved, thus it is relatively easy to identify these *trans* properties in various insects based on their homologies (Carroll et al., 2005). In contrast, *cis* properties appear to be more evolutionarily flexible in a variety of aspects. First, the order of TF binding sites can vary widely within an enhancer region, and the location of enhancers relative to the target gene also appears to be variable. Second, there can be redundancy among multiple enhancers responsible for the same gene (i.e. shadow enhancers) (Hong et al., 2008), allowing them to evolve more rapidly. In addition, the function of each enhancer tends to exhibit low levels of pleiotropy (Carroll, 2008), resulting in the accumulation of more evolutionary changes in enhancers. These characteristics, along with the faster rate of genome evolution in insects compared to vertebrates (Zdobnov and Bork, 2007), make the identification of insect enhancers a challenging task.

Traditionally, enhancers have been identified through reporter assays, in which the transcriptional activation capability of genomic regions near the gene of interest are assessed via a reporter gene construct (see (Suryamohan and Halfon, 2015) to review traditional as well as new methods to identify enhancers). This is a time-consuming and arduous approach, as an enhancer can sometimes reside hundreds of thousands of base pairs away from the gene that the enhancer regulates (Shlyueva et al., 2014). Identification of evolutionarily conserved genomic regions outside of coding regions among several closely related species, such as the *Drosophila* species group, has been helpful in narrowing down regions to survey for enhancer activity (phylogenetic footprinting) (Frazer et al., 2004; Mayor et al., 2000; Sosinsky et al., 2007; Stark et al., 2007). Enhancer predictions based on the TF binding motifs have also been helpful in identifying potential enhancer regions, although the prediction appears to work more efficiently for embryonic enhancers due to the clustering tendency of TF binding motifs within an enhancer active during the syncytial blastoderm stage, while enhancers for other stages might be more difficult to identify through current prediction methods (Li et al., 2007). Combinations of these approaches have allowed for successful identification of enhancers that are active in various developmental contexts in *Drosophila*. More recently, the reporter assay approach has been applied in a genome-wide fashion in *Drosophila* (such as The FlyLight project), identifying over ten thousand genomic regions capable of activating transcription (Jenett et al., 2012; Jory et al., 2012; Kvon et al., 2014; Pfeiffer et al., 2008). Unfortunately, many of these approaches are technically demanding and resource-intensive, and thus are currently only possible in *Drosophila* (but also see (Kazemian et al., 2014) for the successful application of enhancer prediction in non-*Drosophila* insects).

In parallel to the above methods, several genomic approaches have been developed for the identification of possible enhancer regions in the *Drosophila* genome (reviewed in (Shlyueva et al., 2014; Suryamohan and Halfon, 2015)). One such method is Formaldehyde-Assisted Isolation of Regulatory Elements (FAIRE) in combination with next generation sequencing (FAIRE-seq), which identifies open chromatin regions genome-wide (Simon et al., 2012). FAIRE-seq has been used in *Drosophila*, showing that open chromatin regions often correspond to enhancers and other CREs (McKay and Lieb, 2013; Pearson et al., 2016; Uyehara et al., 2017). In addition, FAIRE-seq requires less input material and does not rely on antibodies, thus making it less technically demanding compared to techniques like Chromatin Immunoprecipitation sequencing (ChIP-seq). These features also make FAIRE a promising technique to apply to non-*Drosophila* insects. However, it is important to note that potential enhancers identified by FAIRE (or other genomic approaches) still require functional validations, such as with a reporter assay. This presents another significant hurdle when studying enhancers in non-*Drosophila* insects, as the availability of a modern genetic toolkit (such as a versatile reporter construct) is very limited outside of the *Drosophila* species.

In this study, we set out to establish an enhancer identification and evaluation method in the red flour beetle, *Tribolium castaneum*. *Tribolium* offers a wide variety of genetic and genomic tools, making this insect a powerful model system for comparative developmental biology and evo-devo studies (Denell, 2008; Schmitt-Engel et al., 2015; Tribolium Genome Sequencing et al., 2008; Wang et al., 2007). The robust systemic RNAi response of *Tribolium* has allowed researchers to study *trans* properties in detail (Brown et al., 1999; Bucher et al., 2002; Tomoyasu and Denell, 2004) and identify changes in GRNs responsible for morphological evolution from the *trans* point of view (see (Peel, 2008) to review the findings related to the evolution of insect segmentation; (Tomoyasu et al., 2009) for insect wing evolution; and (Angelini et al., 2012; Smith et al., 2014) for the evolution of insect appendages). However, studies of *cis* properties in *Tribolium* are currently limited due to the lack of reliable enhancer identification methods.

For the initial identification of possible enhancer regions, we first implemented FAIRE-seq and obtained genome-wide chromatin profiles from various tissues and stages of *Tribolium*. The comparison of chromatin profiles between different tissues and stages revealed a distinct set of open chromatin regions in each tissue and stage. Overall, the *Tribolium* open chromatin characteristics are similar to that of *Drosophila*, however, we also noticed some features unique to the *Tribolium* chromatin profiles. Comparison of the FAIRE data to the candidate enhancer regions in the *Tribolium* genome predicated by SCRMshaw (Supervised Cis-Regulatory Module) (Kantorovitz et al., 2009; Kazemian et al., 2011; Kazemian et al., 2014) revealed a very high (>75%) overlap between the two datasets. In addition, we compared the FAIRE profile to the two previously reported enhancer analyses in *Tribolium* (Cande et al., 2009; Eckert et al., 2004; Kazemian et al., 2014; Wolff et al., 1998), and found that the enhancers identified in these studies match well with the open chromatin regions detected by FAIRE.

Second, we chose the wing expression of *nubbin* (*nub*) as a case study, and sought to establish a reporter assay system in *Tribolium*. We initially tested the enhancer activity of the open chromatin regions near the *Tribolium nub* locus in *Drosophila*, and identified a region ~40kb upstream of the *Tribolium nub* gene that has wing enhancer activity in *Drosophila*. This region appears to be open uniquely in the wing tissue, thus providing further support for the ability of FAIRE-seq to identify tissue specific enhancers. In parallel, we also identified the wing enhancer of the *Drosophila nub* gene. Then, we made several reporter constructs and tested these constructs in *Tribolium* using the identified *Drosophila* and *Tribolium nub* wing enhancers. We found that the choice of the core promoter is key in establishing a functional reporter assay system in *Tribolium*, and that a construct with the *Drosophila* Synthetic Core Promoter (DSCP) works well in *Tribolium*. This outcome also shows that the region near the *Tribolium nub* locus with wing enhancer activity in *Drosophila* indeed acts as a wing enhancer in *Tribolium*. In addition, using *hunchback* (*hb*) as another example, we demonstrated that our DSCP reporter construct works in other developmental contexts in *Tribolium*.

Taken together, our results demonstrate that FAIRE-based chromatin profiling by FAIRE-seq, along with the reporter assay system established in this study, is quite powerful at identifying enhancers, and thus will be useful to study the evolution of *cis*-language in *Tribolium*. In addition, our approach might be applicable in other insects for investigating enhancer function and evolution, which will be advantageous for gaining a more comprehensive understanding of the evolution of *cis*-language.

## RESULTS

### FAIRE-seq revealed a spatially and temporally regulated chromatin profile in the *Tribolium* genome

To obtain chromatin profiles from diverse tissues and stages of *Tribolium*, we performed FAIRE-seq with the following six samples; three stages of embryos (0-24 hours, 24-48 hours, and 48-72 hours), the second (T2) and third thoracic (T3) epidermal tissues of the last instar larvae that contain the forewing (elytron) and hindwing imaginal tissues, and the brain isolated from the last instar larvae. The sequence reads obtained from these FAIRE-seq were then mapped to the *Tribolium* genome assembly (Tcas_3.0). Each sample displays a unique set of open chromatin regions (referred to as “peaks”. See Fig. 2A for example), indicating that the FAIRE-seq with the *Tribolium* tissues was successful. The overall open chromatin characteristics are similar between *Tribolium* and *Drosophila*, however, we also noticed some features unique to the *Tribolium* chromatin profiles. We detected more than 40,000 open chromatin regions in the *Tribolium* genome across the samples (Table 1). To identify differences in open chromatin profiles between samples, we performed differential peak calling using DiffBind (FDR < 0.05). The number of differentially accessible peaks between pairs of samples varied widely. For example, there are over 26,000 differentially accessible peaks between 0-24 hours embryos and T3 (Table 1, Fig. S1), reflecting the extensive differences in *cis*-regulatory control that likely exist between these two samples. By contrast, we found only 4 differentially accessible peaks between T2 and T3. The similarity in open chromatin profiles between T2 and T3 tissues is remarkable given the dramatic differences in morphology between forewing and hindwing in *Tribolium*. However, similar findings were obtained in *Drosophila*, suggesting that both species utilize shared sets of enhancers to shape their appendages. Intriguingly, while the level of nucleosome depletion in the FAIRE-isolated genomic regions is variable between stages and tissues, their positions appear to highly correlate with GC-rich regions of the genome (Fig. S2 A). Furthermore, these GC-rich/FAIRE-identified regions occur with a regular interval, producing a “ruler-like” pattern of FAIRE peaks throughout the genome (Fig. S2 B). This regular periodicity of the GC-rich/FAIRE-identified regions appears to be unique to the *Tribolium* lineage, as we did not detect a similar periodicity in other coleopteran genomes or the genome of a lepidopteran, *Bombyx mori* (Fig. S2 C for *Drosophila*. Data not shown for other insects).

**Table 1.** The number of differentially open peaks

|  |  |  |  |  |  |
| --- | --- | --- | --- | --- | --- |
| **T2** | 1 / 3 |  |  |  |  |
| **brain** | 15427 / 7262 | 11634 / 5258 |  |  |  |
| **48-72hr** | 6428 / 8134 | 5575 / 7259 | 1602 / 4089 |  |  |
| **24-48hr** | 10380 / 6729 | 9572 / 6031 | 3162 / 6689 | 863 / 40 |  |
| **0-24hr** | 17450 / 8800 | 14138 / 6808 | 9002 / 8279 | 7651 / 1041 | 2407 / 586 |
|  | **T3** | **T2** | **brain** | **48-72hr** | **24-48hr** |

### Comparison of the FAIRE data to previous enhancer studies in *Tribolium*

There are several previous works investigating the activity of *Tribolium* enhancers. To our knowledge, Eckert et al. is the only study analyzing enhancer activity in the *Tribolium* native context, which identified enhancers important for the stripe expression of the *Tribolium hairy* gene (Eckert et al., 2004). Some additional enhancers for *Tribolium* genes have also been identified, albeit in a cross-species context (i.e. *Drosophila*). These include enhancers for *hunchback* (Wolff et al., 1998), *single-minded*, *cactus* and *short gastrulation* (Cande et al., 2009), *labial*, *Dichaete*, and *wingless* (Kazemian et al., 2014). We analyzed the FAIRE profiles at these gene loci and found that FAIRE peaks match with many of the enhancer regions identified through these studies (Fig. S3).

More recently, Kazemian et al. applied their enhancer discovery approach, SCRMshaw, to the *Tribolium* genome and predicted 1214 genomic regions as potential enhancers (Kantorovitz et al., 2009; Kazemian et al., 2014). Comparison of the FAIRE data to the SCRMshaw predictions reveals a striking degree of overlap between the two datasets: 78.8% (957/1214) of SCRMshaw predictions overlap at least one embryonic FAIRE peak, while 88.1% (1070/1214) of predictions overlap at least one larval FAIRE peak (Table. S1, S2; *P* ≈ 0); overall, 1096 of the 1214 predicted CRMs (90.3%) overlap at least one FAIRE peak. For certain sets of SCRMshaw predictions, the overlaps are even more extensive: for example, 98% (97/99) of wing-specific predicted CREs overlap a larval FAIRE peak (Table. S1). Taken together, the high degree of overlap between the FAIRE peaks and previously identified enhancer regions, and FAIRE-peaks and SCRMshaw-predicted CREs, verifies that FAIRE-seq is a powerful tool to identify enhancers in *Tribolium*.

### Identification of the *Tribolium nub* wing enhancer using an inter-species reporter assay

As mentioned in the introduction, reporter assays are a time consuming and laborious task, which makes it difficult to perform in non-*Drosophila* insects, including *Tribolium*. However, to be able to fully exploit the benefit of the FAIRE profiling data, it will be critical to have a reliable method to evaluate the function of *Tribolium* enhancers. The activity of some potential *Tribolium* enhancers has been successfully evaluated via a reporter assay in *Drosophila* (Cande et al., 2009; Kazemian et al., 2014; Wolff et al., 1998; Zinzen et al., 2006). We reasoned that the enhancer of a gene that has a conserved expression pattern (both temporal and spatial) between *Drosophila* and *Tribolium* has the highest chance of being active, even in an inter-species context, and is thus ideal for a case study. The enhancer responsible for the wing expression of *nub* fits this criterion, as *nub* is expressed broadly in the tissues that give rise to wings in both insects (Fig. 1) (Ng et al., 1995; Tomoyasu et al., 2009). In addition, there is an enhancer trap line for *nub* available in *Tribolium* (*pu11*. Fig. 1C, D). We have previously determined that this enhancer trap line has a piggyBac construct inserted about 30kb upstream of the *nub* transcription start site (Clark-Hachtel et al., 2013), which can be used as a starting point to survey for the wing enhancer.

**Fig. 1.**
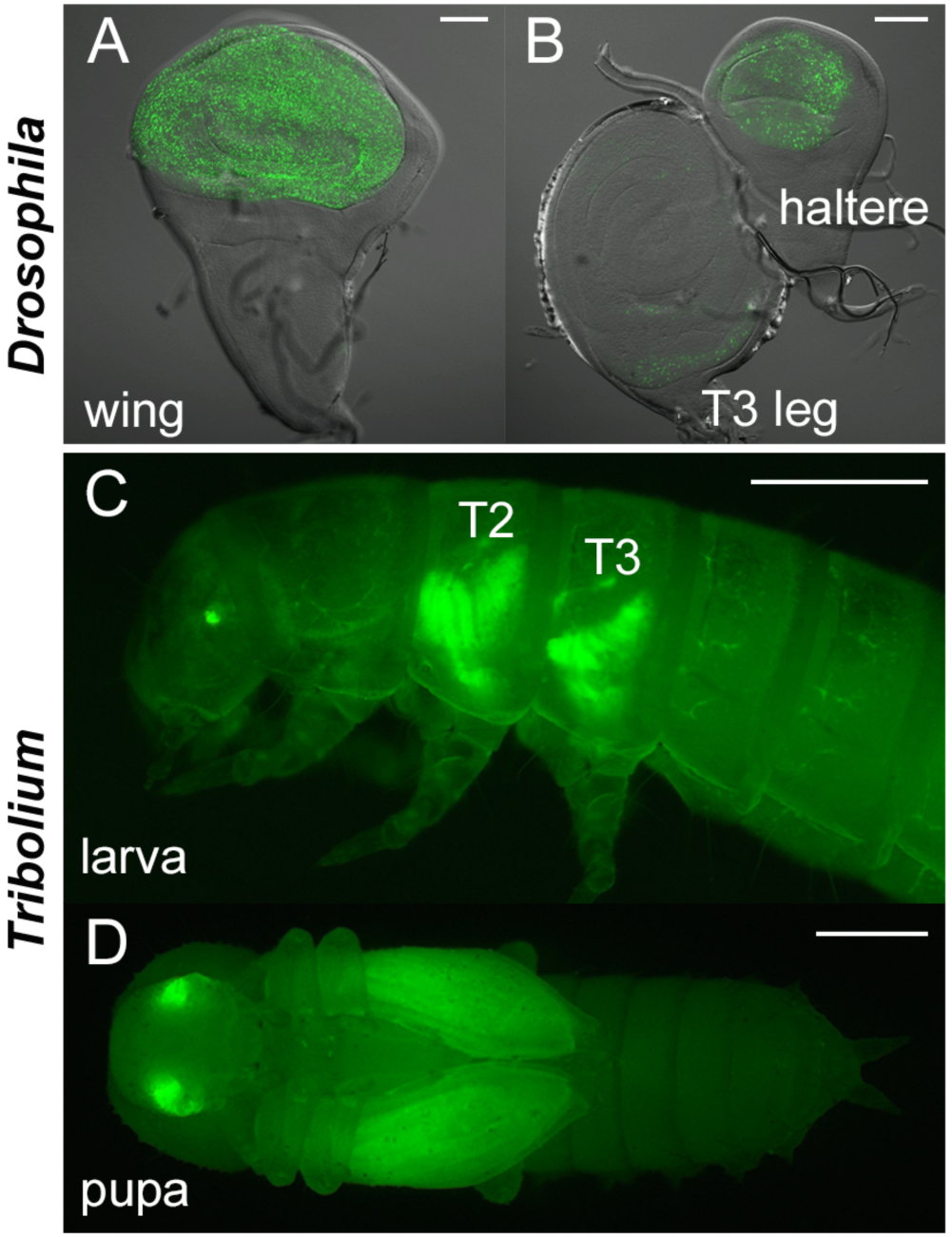
*nub* enhancer trap expression in *Drosophila* and *Tribolium*. (A, B) The *nub* enhancer trap expression in the wing disc (A), and the haltere and T3 leg discs (B) in *Drosophila*. (C, D) Expression pattern of the *nub* enhancer trap line (*pu11*) at the larval (C) and pupal (D) stage in *Tribolium*. Scale bar: 0.5mm.

*nub* codes for an evolutionarily conserved transcription factor important for the proliferation of wing cells (Ng et al., 1995). *Drosophila* has two *nub* paralogs (*nub* and *pdm2*), while *Tribolium* has one (*Tc-nub*). FAIRE analysis has revealed a number of open chromatin regions located in and near the *Tc-nub* locus (Fig. 2A). Some of the open chromatin regions are shared across the six samples tested (such as the region corresponding to the promoter), but others are unique to specific tissues and stages. We decided to test the two open chromatin regions at or near the *pu11* insertion site (*Tc-nub3* and *Tc-nub2*) in *Drosophila* (Fig. 2A, B). In addition, we also tested another major open chromatin region located further upstream of the *pu11* insertion site (*Tc-nub1*). This region is open predominantly in the larval T2 and T3 epidermal tissues (containing the future wing tissues) but not in any of the embryonic samples, suggesting the possibility that this region contains enhancers specific to the post-embryonic stage (Fig. 2A, B).

**Fig. 2.**
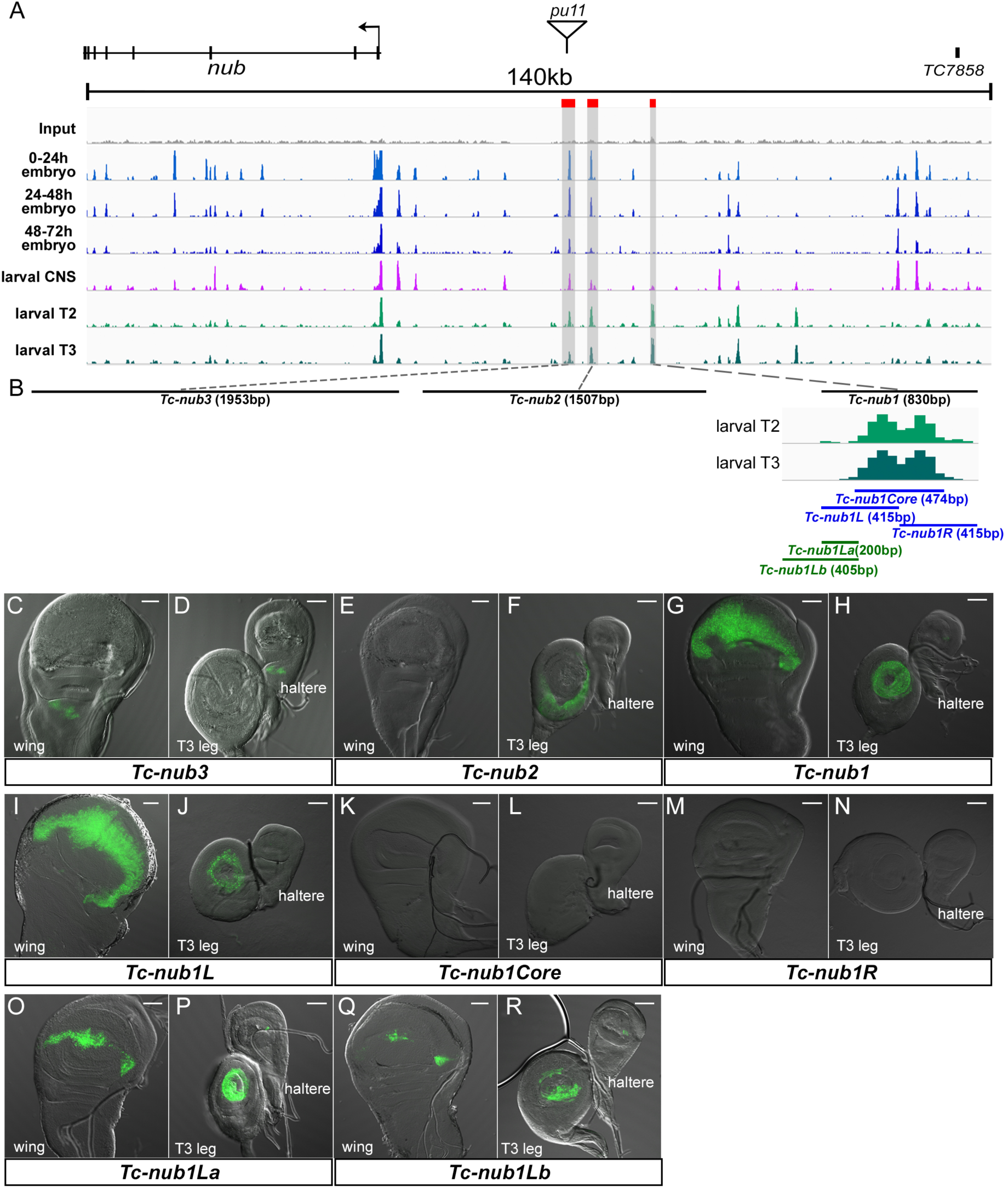
Identification of the *Tribolium nub* wing enhancer with FAIRE and inter-species reporter assay. (A) FAIRE profiles at the *Tribolium nub* locus in six different tissues/stages. The *pu11* insertion site is indicated with a triangle. Three peaks near the *pu11* insertion site chosen for evaluating enhancer activity were marked with red boxes. (B) Summary of the regions tested by the reporter assay. The distance between *Tc-nub1, 2,* and *3* are not scaled. The magnified view of the FAIRE peak corresponding to *Tc-nubL1* is also presented. (C-R) Enhancer activity of each *Tribolium* genomic region tested in the *Drosophila* imaginal discs. Scale bar: 50 µm.

The inter-species reporter assay showed that *Tc-nub2* and *Tc-nub3* do not have enhancer activity in the future wing-related tissues (wing and haltere imaginal discs) when tested in *Drosophila* (Fig. 2C-F). *Tc-nub*3 showed activity in a small region near the hinge of the wing and haltere disc, but not in the region that gives rise to the wings (wing and haltere pouches) (Fig. 2C, D). *Tc-nub*2 drove the reporter expression in the leg discs, but did not show any enhancer activity in the wing and haltere discs (Fig. 2E, F). In contrast, *Tc-nub1* showed significant enhancer activity in the pouch region of the wing disc (Fig. 2G). *Tc-nub1* also drove reporter expression in the leg disc, but was not active in the haltere disc (Fig. 2H). Since *Tc-nub1* corresponds to the region uniquely open in the larval epidermis in *Tribolium*, the outcome of our inter-species reporter assay indicates that (i) the open chromatin profiling of various tissues and stages by FAIRE-seq in *Tribolium* can help predict tissue/stage specific enhancers from the *Tribolium* genome, and (ii) the inter-species reporter assay can be useful, at least for the enhancers responsible for the post-embryonic expression of *nub* in *Tribolium*.

We next sought to minimize the *Tc-nub* wing enhancer by testing three shorter fragments within the *Tc-nub1* region (Fig. 2B). Interestingly, despite covering the main FAIRE peak region of *Tc-nub*1, *Tc-nub1Core* did not show any enhancer activity in the wing (Fig. 2K, L). Instead, *Tc-nub1L*, which corresponds to only a part of the major open chromatin region, drove the reporter expression with a pattern and level almost identical to *Tc-nub1* (Fig. 2I, J). *Tc-nub1R* did not show any enhancer activity (Fig. 2M, N). These results suggest that the important elements for driving wing expression reside within the first 200bp of *Tc-nub*1. We tested this idea by making a reporter construct using only the 200bp region unique to *Tc-nub1L* (*Tc-nub1La*, Fig. 2B). This fragment drove the reporter expression in the wing and leg discs, albeit with a more restricted expression domain and/or a lower expression level compared to *Tc-nub1L* (Fig. 2O, P). We also tested a construct that contains the *Tc-nub1L* region along with an additional 200bp sequence outside of *Tc-nub1* (*Tc-nub1Lb* in Fig. 2B), since the location of the functional *Tc-nub* wing enhancer may be slightly misaligned with the open chromatin region predicted by FAIRE. However, *Tc-nub1Lb* showed even weaker enhancer activity (Fig. 2Q, R), suggesting that there might be a suppressor element next to the *Tc-nub1* region. The constructs we made also drove reporter expression outside of the wing and leg imaginal disc. These results are summarized in Table S3.

### Identification of the *Drosophila nub* wing enhancer using a combination of genomic resources, FAIRE profiling, and the reporter assay approach in *Drosophila*

As a comparison to the enhancer identified via an inter-species reporter assay described above, we sought to identify the *nub* wing enhancer native to the species used for the reporter assay (i.e. *Drosophila*). As mentioned, there are two *nub* paralogs in *Drosophila* (*nub* and *pdm2*), both of which have similar expression in the wing pouch (Ng et al., 1995). We first took advantage of the FlyLight project and surveyed the *nub* and *pdm2* loci for a genomic region that has wing enhancer activity. Among the 33 constructs tested in FlyLight (Fig. 3A), one region (GMR11F02) has a record of enhancer activity in the wing and haltere pouch, along with additional expression in the leg disc (Fig. 3B, C). We then utilized the previously published FAIRE profile for *Drosophila* (McKay and Lieb, 2013), and identified three distinct regions within GMR11F02 that are open in the wing and haltere discs (Fig. 3A). We cloned these three regions (Fig. 3B, *Dm-nub1*, *Dm-nub2*, and *Dm-nub3*), and tested their enhancer activity in *Drosophila*. Among the three regions, *Dm-nub2* displayed strong enhancer activity in the wing pouch region (Fig. 3G, H). *Dm-nub1* (Fig3E, F) and *Dm-nub3* (Fig3I, J) did not drive reporter expression in the wing and haltere discs. In addition, *Dm-nub3* was active in the leg disc, suggesting that the *nub* wing/haltere enhancer and the leg enhancer are separable (Fig. 3J). To further minimize the *Dm-nub* wing enhancer, we tested three shorter fragments within *Dm-nub2* (*Dm-nub2a*, *Dm-nub2b*, and *Dm-nub2c*. Fig. 3D). The wing related expression is driven by *Dm-nub2a*, albeit at a weaker level (Fig. 3K, L). This suggests that, although *Dm-nub2a* contains sufficient components to drive wing expression, a broader genomic region is required for robust wing expression of *Dm-nub*. In contrast, *Dm-nub2b* and *Dm-nub2c* did not drive any expression (Fig. 3M-P). The expression patterns of these constructs in other tissues are summarized in Table S4. Taken together, the *Dm-nub2* region we isolated (1.3kb) is sufficient to drive a robust wing expression in *Drosophila*.

**Fig. 3.**
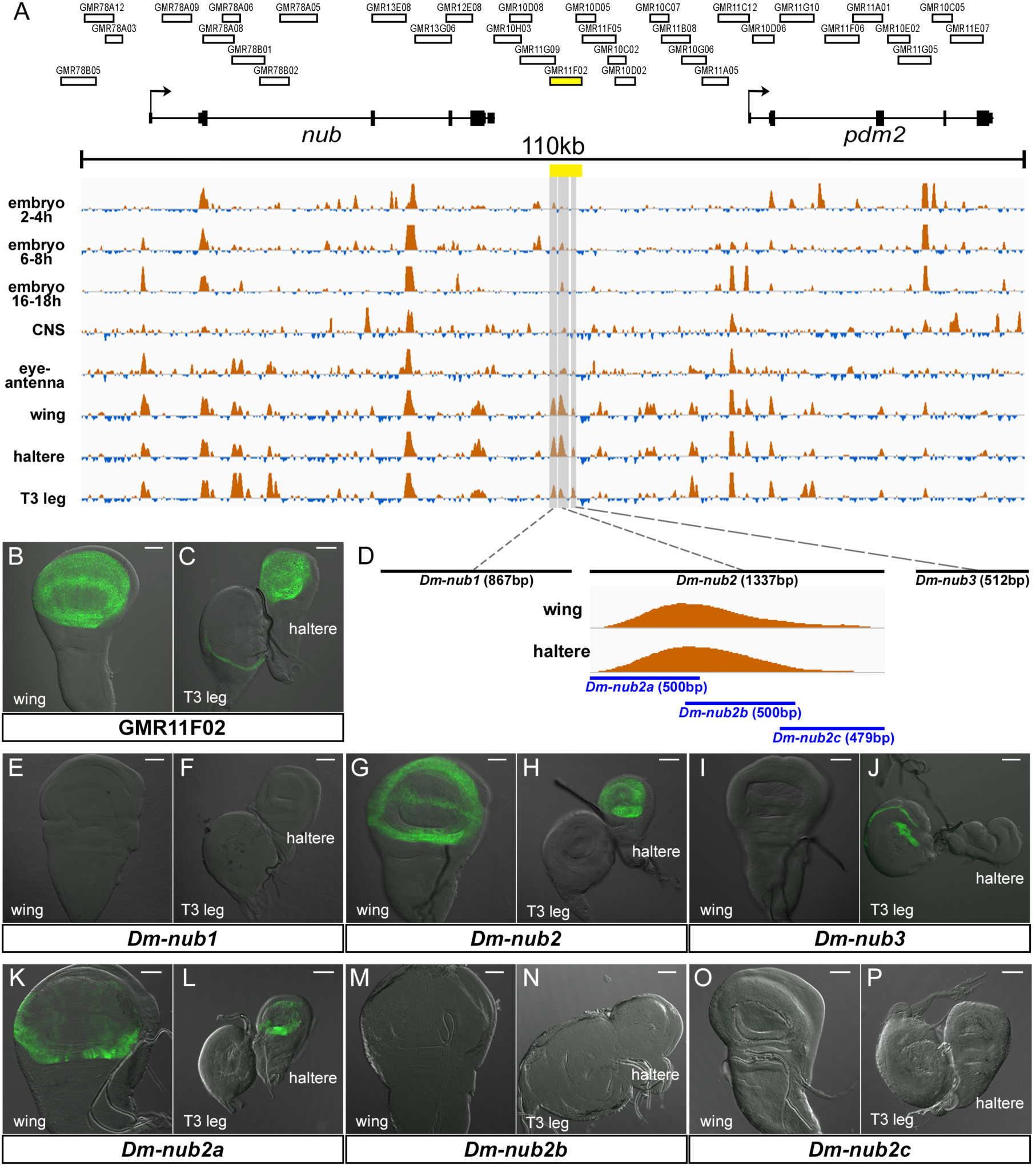
Identification of the *Drosophila nub* wing enhancer. (A) FAIRE profiles from eight different tissues/stages at the *nub* and *pdm2* loci in *Drosophila*. The regions surveyed in the FlyLight project are also indicated. The region that shows wing enhancer activity is marked by yellow. (B, C) Expression driven by GMR11F02 in the *Drosophila* imaginal discs. (D) Summary of the regions within GMR11F02 tested by the reporter assay. The relative distance between *Dm-nub1, 2,* and *3* are not scaled. The magnified view of the *Dm-nub2* peak is also included. (E-P) Enhancer activity of each *Drosophila* genomic region tested in the *Drosophila* imaginal discs. Scale bar: 50 µm.

### Establishing a reporter assay system and evaluating the *nub* wing enhancers in *Tribolium*

Although some *Tribolium* enhancers were shown to be active in the cross-species context, these enhancers still need to be examined in their native species for functional validation. However, the lack of a reliable reporter construct has been a major obstacle in performing functional evaluation of enhancers in *Tribolium*. The GATEWAY system (Katzen, 2007) has been useful in quickly cloning genomic regions into a reporter construct and testing their enhancer activity in *Drosophila*. We sought to establish a GATEWAY compatible reporter construct that is functional in *Tribolium*.

A key issue in establishing a reporter construct is the choice of promoters. Previous studies have raised concerns about using *Drosophila* promoters in *Tribolium* (Schinko et al., 2010). While establishing the Gal4/UAS system in *Tribolium*, Schinko et al. found that the core promoter isolated from a *Tribolium* endogenous gene, *Tc-hsp68*, worked more efficiently when compared to the exogenous promoters that were tested (Schinko et al., 2010). We therefore made a GATEWAY compatible piggyBac construct that contains the *Tc-hsp68* core promoter driving the *dsRed* gene (piggyGHR, Fig. 4A). In addition, we added the *gypsy* element, a *Drosophila* insulator, flanking the reporter assay construct to prevent position effects (Fig. 4A). We tested this piggyBac construct with the *Tribolium* and *Drosophila nub* wing enhancers (*Tc-nub1L* and *Dm-nub2*) in *Drosophila*. Both *Tc-nub1L* and *Dm-nub2* drove dsRed expression identical to the patterns obtained with the *Drosophila* reporter construct (compare Fig. 4B, C to Fig. 2I, J, and Fig. 4D, E to Fig. 3G, H), confirming that piggyGHR is functional. However, neither *Tc-nub1L* or *Dm-nub2* in piggyGHR showed consistent enhancer activity in the wing tissues when transformed into *Tribolium* (Fig. 4F-M). Among the seven independent transgenic lines obtained for piggyGHR*-Tc-nub1L*, none of them had clear dsRed expression in the wing tissues (Fig. 4F-K). Instead, four lines had dsRed expression in non-wing tissues, with a distinct pattern in each line, likely due to trapping local enhancers (Fig. 4F-K). We obtained only two independent transgenic lines for piggyGHR-*Dm-nub*2, neither of which had dsRed expression in the wing tissue (Fig. 4L, M). These results indicate that our construct with the *Tc-hsp68* core promoter does not work well for reporter assays, at least in the wing related tissues in *Tribolium*, although it does work in *Drosophila*. Alternatively, it is also possible that the *Drosophila gypsy* insulators we added to the construct might not be functioning properly in *Tribolium*.

**Fig. 4.**
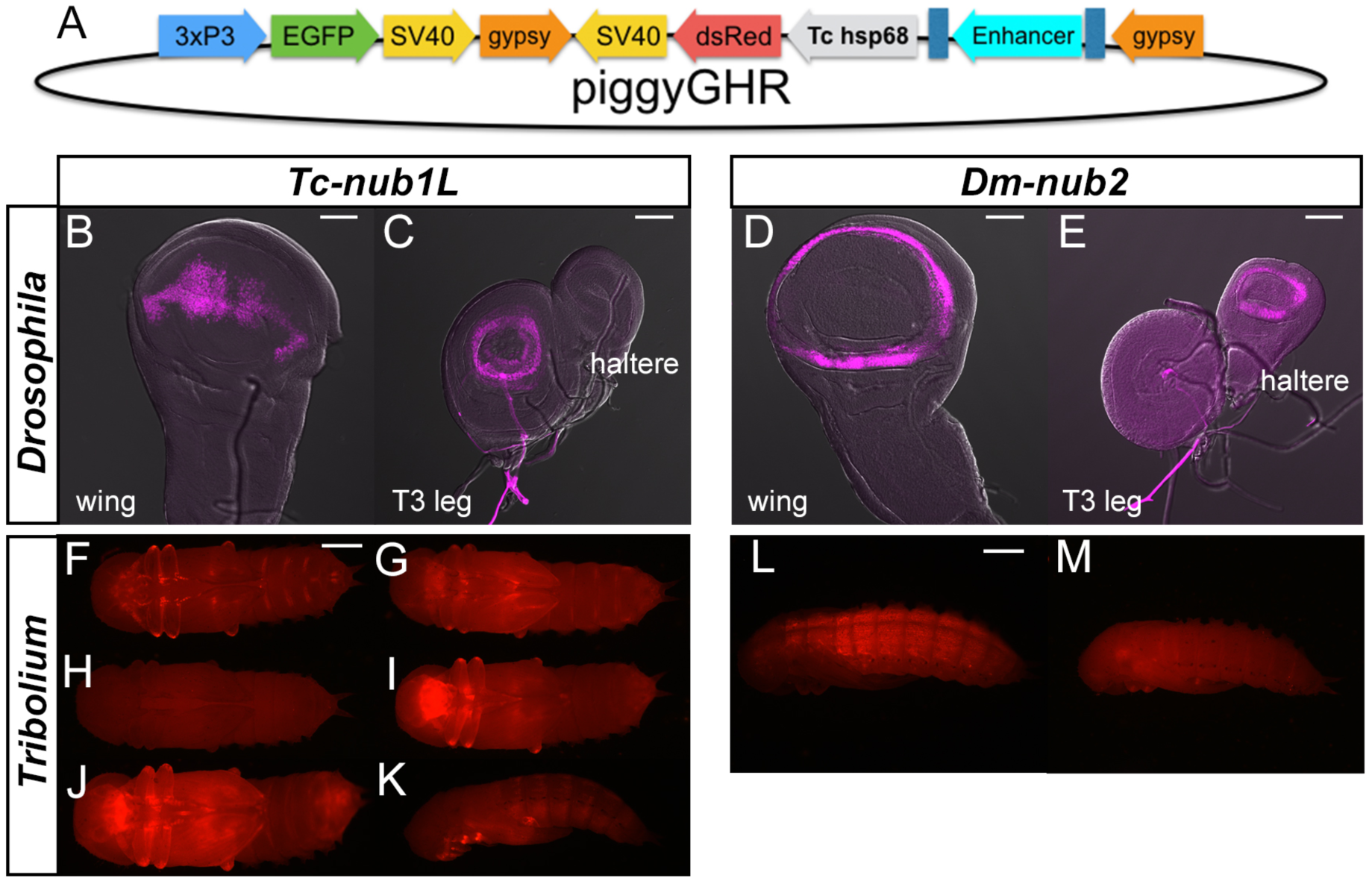
Reporter assay with the *Tc-hsp68* promoter construct in *Drosophila* and *Tribolium*. (A) The piggyGHR construct. (B-E) Enhancer activity of *Tc-Nub1L* (B, C) and *Dm-nub2* (D, E) tested with the piggyGHR construct in *Drosophila*. (F-M) Enhancer activity of *Tc-nub1L* (F-K) and *Dm-nub2* (L, M) tested with piggyGHR at the pupal stage in *Tribolium*. Six independent lines for *Tc-nub1L* (F-K) and two for *Dm-nub2* (L, M) were shown. Scale bar: 50 µm (B-E), 0.5 mm (F-M)

We next tested a synthetic promoter in *Tribolium*. Pfeiffer et al. modified the Super Core Promoter 1 (SCP1) (Juven-Gershon et al., 2006) and constructed the *Drosophila* Synthetic Core Promoter (DSCP), which was used for the FlyLight project as well as other *Drosophila* reporter constructs including pFUGG used in this study (McKay and Lieb, 2013). We made a piggyBac construct with the DSCP driving mCherry (piggyGUM, Fig. 5A). We decided to remove the *Drosophila gypsy* insulators from our construct to avoid possible inter-species issues. Similar to piggyGHR, piggyGUM with the *Drosophila* and *Tribolium nub* wing enhancers drove reporter expression in the wing disc in *Drosophila* (Fig. 5B-E), confirming that piggyGUM is functional. In *Tribolium*, in contrast to the piggyGHR constructs, piggyGUM-*Tc-nub1L* successfully recaptured the expression pattern of the *nub* enhancer trap line (*pu11*) and drove reporter expression in the wing related tissues (both in T2 and T3) at both larval and pupal stages (Fig. 5F-I, compared to Fig. 1C, D). piggyGUM-*Dm-nub2* also showed enhancer activity in the larval wing discs in *Tribolium* (Fig. 5J-L). The expression driven by *Dm-nub2* in *Tribolium* was mostly in the wing hinge and the margin regions, similar to the pattern observed for this enhancer in the *Drosophila* imaginal discs (Fig. 3G, H, Fig. 5D, E). These results indicate that (i) our GATEWAY compatible DSCP piggyBac construct (piggyGUM) can be used for reporter assays both in *Tribolium* and *Drosophila*, and (ii) the *Tribolium nub* wing enhancer identified through an inter-species reporter assay (*Tc-nub1L*) is indeed functional as a wing enhancer in *Tribolium*. It is also worth mentioning that some of the piggyGUM transgenic lines showed mCherry expression in tissues outside of wings (data not shown). The expression patterns outside of the wing related tissues were not consistent among the transgenic lines, suggesting that the piggyGUM construct also occasionally traps endogenous enhancers.

**Fig. 5.**
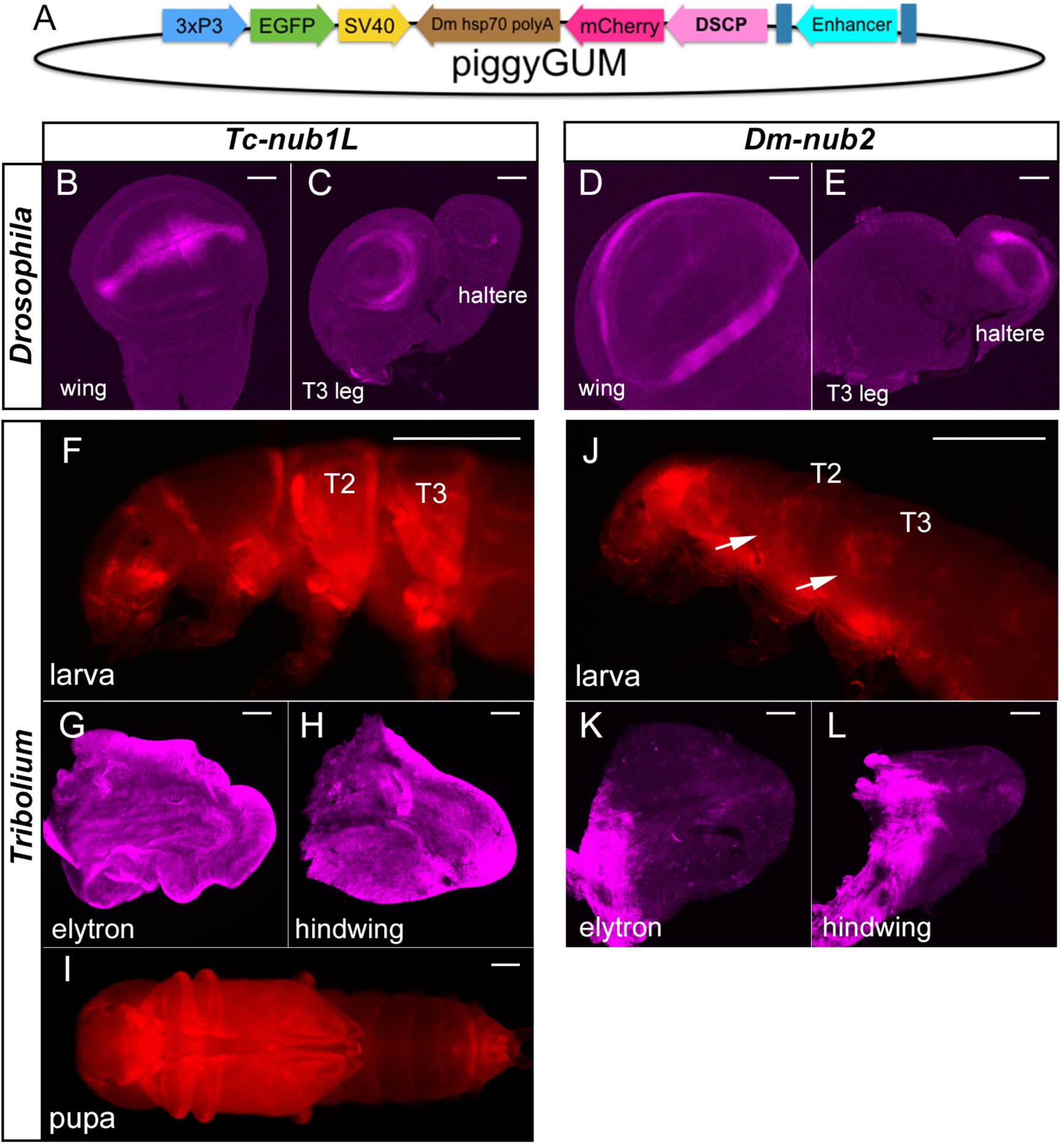
Reporter assay with the *Drosophila* synthetic core promoter construct in *Drosophila* and *Tribolium*. (A) The piggyGUM construct. (B-E) Enhancer activity of *Tc-Nub1L* (B, C) and *Dm-nub2* (D, E) tested with the piggyGUM construct in *Drosophila.* (F-L) Reporter expression of piggyGUM-*Tc-nub1L* (F-I) and piggyGUM-*Dm-nub2* (J-L) in *Tribolium*. Scale bar: 50 µm (B-E, G, H, K, L), 0.5 mm (F, J, I)

We also tested if the promoter endogenous to the enhancer works better for a reporter assay construct in *Tribolium*. We made a piggyBac construct with the 2kb sequence upstream of the *Tc-nub* transcription start site (confirmed by 5’ RACE (Clark-Hachtel et al., 2013)) as the promoter (Fig. 6A, piggyNub-proR). We also used the 2kb downstream of the *Tc-nub* stop codon (confirmed by 3’ RACE (Clark-Hachtel et al., 2013)) as the 3’ untranslated region (UTR) and the poly A signal native to *Tc-nub* (Fig. 6A). We made a similar construct for Tc-Act5c (1kb upstream of the transcription start site and 1kb downstream of the stop codon as the native promoter and polyA signal, respectively) as a comparison (Fig. 6B). To our surprise, *Tc-nub1L* in piggyNub-proR did not drive any expression in *Tribolium* (Fig. 6C-F) or in *Drosophila* (Fig. 6G, H). Realtime-qPCR analysis revealed that there is no transcription of dsRed in these transgenic lines in both species (data not shown), suggesting that the lack of reporter expression is not due to incompatibility of the reporter gene with the *Tc-nub* UTRs and is rather due to the *nub* wing enhancer failing to work with the endogenous promoter and/or polyA signal. In contrast to piggyNub-proR-*Tc-nub1L*, piggyAct5cR shows strong and ubiquitous dsRed expression in *Tribolium* (Fig. 6I), indicating that our strategy of incorporating the endogenous transcription and translation components is valid. Intriguingly, however, piggyAct5cR did not drive any expression in *Drosophila* (data not shown), implying a strict species specific nature of the transcription and/or translation components (such as promoters), even for an evolutionarily highly conserved house-keeping gene that is uniformly expressed in various species including *Drosophila* and *Tribolium* (Chung and Keller, 1990).

**Fig. 6.**
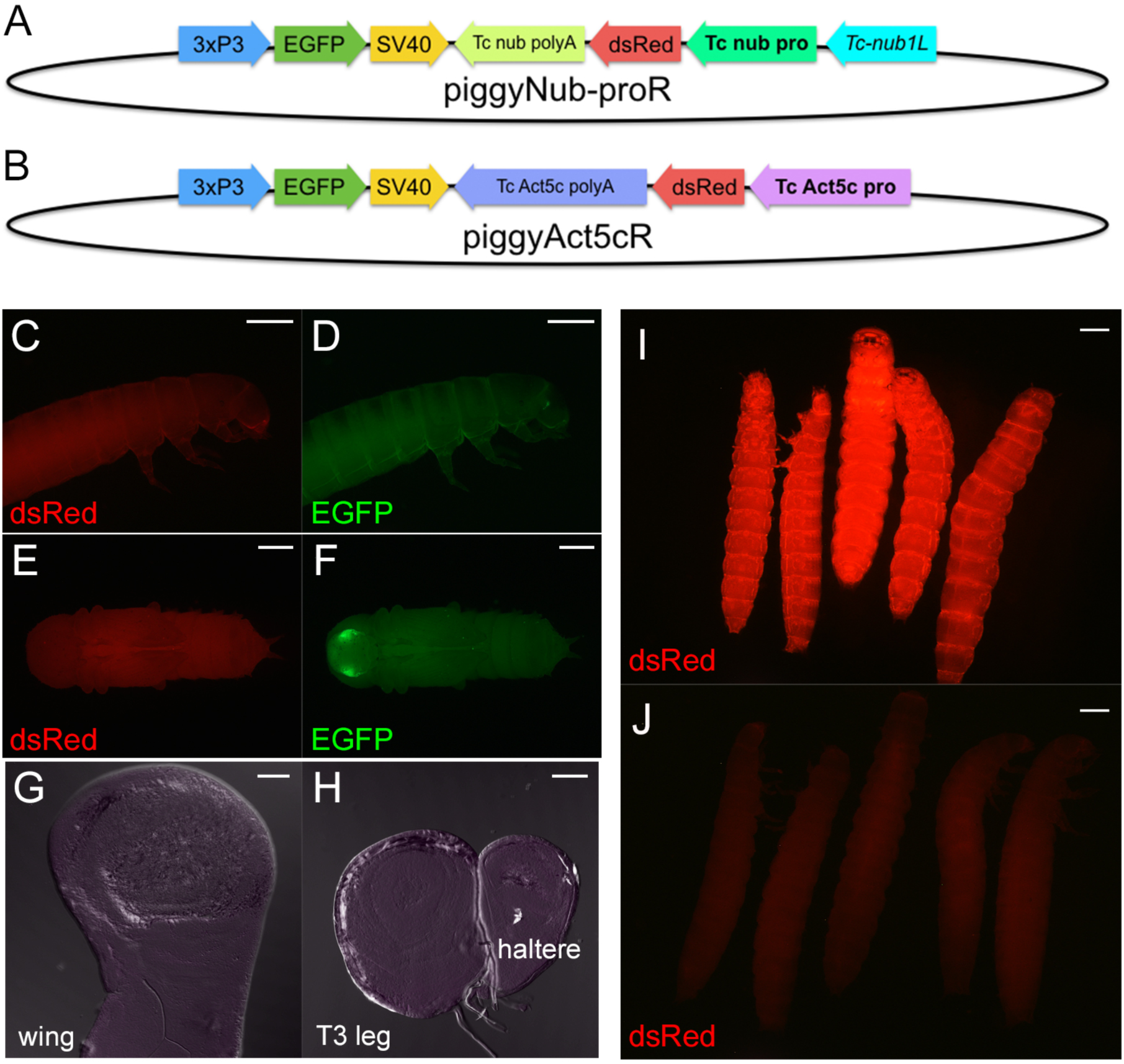
Reporter assay with the *Tribolium* endogenous promoters in *Drosophila* and *Tribolium.* (A) The piggyNub-proR construct. (B) The piggyAct5cR construct. (C-F) Enhancer activity of *Tc-Nub1L* tested with the piggyNub-proR construct. dsRed reporter expression is completely absent (C, E), even though EGFP (D, F) confirms the presence of the construct transgened. (G, H) The piggyNub-proR reporter expression in *Drosophila* imaginal discs. (I) dsRed expression of the piggyAct5cR at the larval stage in *Tribolium*. (J) dsRed expression of the piggyNub-proR with *Tc-Nub1L* at the larval stage in *Tribolium*, with the same exposure time as (I). Scale bar: 0.5 mm (C-F, I, J), 50 µm (G, H).

### Testing the reporter construct in another context in *Tribolium*

We next wondered if our DSCP reporter system works in a context other than wings in *Tribolium*. We chose *hb* as a case study, and tested the reporter activity during embryogenesis. *hb* expression in *Tribolium* starts as a broad posterior domain in the blastoderm and clears from posterior to form an anterior band of expression that covers pre-gnathal and gnathal segments (Lynch et al., 2012; Marques-Souza et al., 2008). In the early germband stage, the band resolves into a stripe covering the labium (Fig. 7B) (Marques-Souza, 2007; Zhu et al., 2017). Wolff et al. previously identified a genomic region at the *Tribolium hb* locus that drives blastoderm expression when introduced in *Drosophila* (Fig. 7A, orange bar) (Wolff et al., 1998). This region corresponds to a SCRMshaw prediction (Fig. 7A, purple bars). Therefore, although the FAIRE signal at this region is weak (likely due to the wide time window of sampling during early embryogenesis), the outcomes of previous studies make this region an excellent candidate enhancer to test with our reporter system in *Tribolium*. We cloned a 1340bp fragment containing this genomic region (*hb*-PE1, Fig. 7A, red bar), and tested its enhancer activity using the piggyGUM construct in *Tribolium*. *in situ* hybridization for the *mCherry* reporter gene revealed that the piggyGUM-PE1 construct recapitulates the *hb* expression at the early germband stage in *Tribolium* (Fig. 7C). This result indicates that (i) our DSCP reporter system works well even during embryogenesis in *Tribolium*, and (ii) *hb*-PE1 contains the *hb* early germband enhancer.

**Fig. 7.**
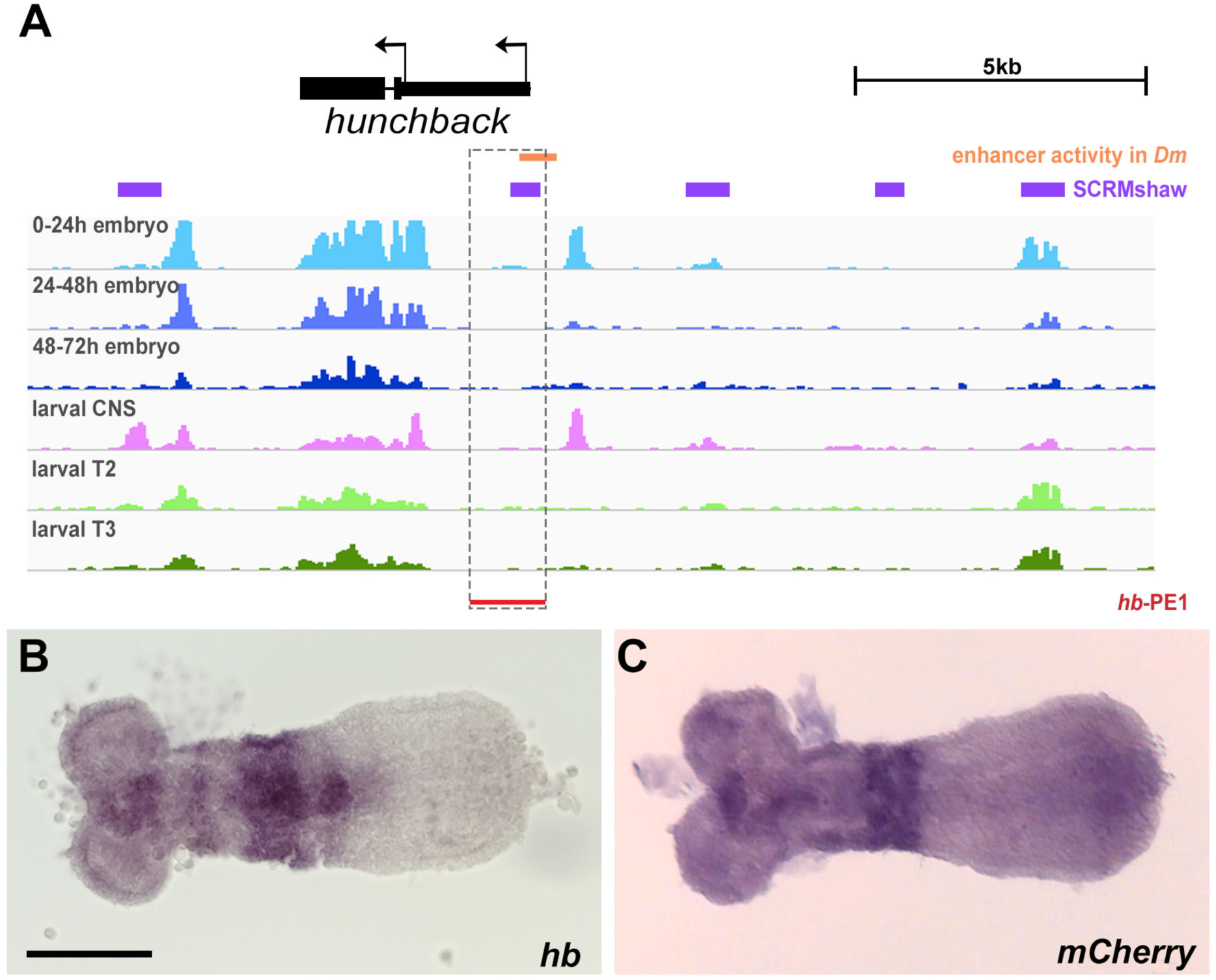
*hb* enhancer analysis in *Tribolium*. (A) FAIRE profiles at the *hb* locus. Orange bar: blastoderm enhancer activity when introduced in *Drosophila*, purple bars: SCRMshaw predictions, red bar: the 1340bp fragment tested in this study (*hb*-PE1). (B) *hb* expression at the early germband stage detected by *in situ* hybridization for *hb* transcript. (C) *mCherry* reporter gene expression of piggyGUM-*hb*-PE1 detected by *in situ* hybridization for *mCherry* transcript. Scale bar: 100 µm (B, C).

In summary, we established a functional reporter assay system that works in diverse developmental contexts in *Tribolium* and also successfully identified the enhancers responsible for wing expression of *nub* and early germband expression of *hb*. Furthermore, our reporter construct (piggyGUM) is compatible in both *Drosophila* and *Tribolium*, implying that this reporter construct may be applicable even to other insect species.

## DISCUSSION

In this study, we demonstrated that FAIRE-based chromatin profiling is a powerful approach for identifying CREs, such as enhancers, in *Tribolium*. The *Tribolium nub* wing enhancer we identified (*Tc-nub1L*) is over 40kb away from the *nub* transcription start site, and 10kb away from the *pu11* insertion site, which would be very difficult to identify without the aid of open chromatin profiles. In addition, with the usage of the DSCP, we were able to establish a functional reporter assay construct in *Tribolium*. Combination of FAIRE-based chromatin profiling with this reporter assay system will allow us to assess the function and evolution of enhancers in *Tribolium*.

### FAIRE profiles in *Tribolium*

Genome-wide FAIRE profiling in *Tribolium* has identified a significant number of genomic regions whose chromatin status is regulated in a tissue and stage specific manner (Table 1, Fig. S1). These regions are promising candidates for future enhancer studies in *Tribolium*. In addition, our FAIRE analysis has revealed both evolutionarily conserved and diverged aspects of chromatin state regulation between *Drosophila* and *Tribolium*. For the conserved aspect, we saw similar chromatin profiles for the T2 and T3 epidermal samples, even though these two tissues differentiate into morphologically distinct structures (the elytron in T2 and hindwing in T3). This outcome echoes the message obtained from the *Drosophila* FAIRE study, namely that chromatin profiles are largely similar among the similar lineages of tissues (such as legs, wings, and halteres), with the exception of a handful of “master control gene” loci (McKay and Lieb, 2013). In fact, three of the four differentially-open FAIRE peaks between T2 and T3 in our *Tribolium* FARE analysis are within the *Ultrabithorax* (the T3 selector gene) locus (Fig. S1) (to review the function of *Ultrabithorax* during wing development, see (Tomoyasu, 2017)). In contrast, the *Tribolium* FAIRE profiles during embryogenesis show an interesting difference when compared to those in *Drosophila*. In *Drosophila*, the number of genomic regions that are open is fairly consistent throughout embryogenesis, with a distinct set of genomic regions being open in each stage (McKay and Lieb, 2013). In *Tribolium*, we noticed that a larger number of chromatin regions are open early in embryogenesis, and some of these regions are subsequently closed, resulting in a smaller number of open chromatin regions in later stages. This difference may be a reflection of the different modes of embryogenesis found in the two insects (long vs. short germ band embryogenesis), although the significance of the difference in chromatin profiles is yet to be investigated.

We also noticed a strict overlap between the GC-rich regions and FAIRE-detected open chromatin regions. This raises an interesting argument about the evolution of enhancers. Are these regions open because they are functionally important (such as enhancers)? Or, have these regions become enhancers, because they were open due to a bias in their nucleotide content and thus accessible to transcription factors? There appears to be a similar correlation among the GC-rich regions, enhancers, and FAIRE peaks in *Drosophila* (Li et al., 2007; McKay and Lieb, 2013). It will be interesting to investigate how GC-rich regions overlap with open chromatin regions in other insects. In addition, we found that the GC-rich/FAIRE-positive regions appear in a regular interval throughout the *Tribolium* genome. The molecular basis and functional implication of this periodicity is currently unknown, however, it is intriguing to speculate that a genome-wide event (such as transposon invasion) might have significantly influenced the chromatin state landscape in the *Tribolium* lineage.

### Overlaps between FAIRE and SCRMshaw enhancer prediction

The high degree of overlap observed between FAIRE peaks and enhancers predicted by the completely different, solely computational, SCRMshaw method provides further confirmation that FAIRE is an effective means for enhancer discovery in *Tribolium*. Overall, the number of FAIRE peaks is well in excess of the number of SCRMshaw predictions. Several factors likely account for this result. First, the SCRMshaw predictions were performed at high stringency in order to minimize potential false-positive results (Kazemian et al., 2014); relaxing the prediction criteria would yield more predicted enhancers. While this would potentially lead to more false positives, the >90% overlap seen for several specific data sets (Table S1) suggests that stringency could be relaxed in at least some cases. Second, SCRMshaw relies on training data from known *Drosophila* enhancers; therefore enhancers with properties significantly deviating from those of *Drosophila* enhancers will be found only by chromatin profiling, such as FAIRE. Finally, although FAIRE appears to be biased toward enhancers (Song et al., 2011), it also identifies other regions of open chromatin such as promoters and insulator regions (Giresi et al., 2007), which are not predicted by the enhancer-specific SCRMshaw.

The twin issues of higher SCRMshaw false-positive rates at lower prediction stringencies and FAIRE’s lack of discrimination with respect to enhancers with specific spatial and temporal activity profiles suggest that considerable advantages could be obtained by using the methods in combination. Overlap with FAIRE peaks can be used to filter out false-positive SCRMshaw predictions, allowing predictions to be performed at lower stringency and thus higher sensitivity. Conversely, SCRMshaw prediction can be used to focus on potentially more relevant FAIRE peaks, helping to avoid selecting sequences representing enhancers active in tissues other than the one of interest; enhancers for a neighboring housekeeping gene; insulators; and cryptic promoters or those for unannotated genes. This will be particularly useful for situations like the one seen here for the larval samples, where cleanly separating wing from body wall tissue was difficult, a common challenge when attempting to isolate tissues from small organisms such as insect embryos.

### Enhancer activity in inter-species contexts and the limitation of non-native reporter assays

Our reporter assays in two insect species showed that both *Drosophila* and *Tribolium nub* wing enhancers were at least partially active in the inter-species context. We identified a 20-bp sequence shared between the two enhancers that contains binding sites of some wing-related transcription factors (such as Brinker and Mad) (Fig. S4). However, deletion of this sequence did not influence the activity of these enhancers when tested in *Drosophila*, indicating that this sequence is dispensable for enhancer function (Fig. S4). We did not recognize any other significant sequence similarity or a conserved TF-binding site architecture between the two enhancers, suggesting that the regulatory landscape in the wing of the two species is evolutionarily maintained (as the *nub* enhancers can be functional in inter-species contexts) despite the lack of noticeable sequence conservation in the enhancer itself. A thorough examination of *trans* properties that regulate the *nub* wing enhancers may give us insights into how enhancers evolve under a conserved regulatory landscape.

Although the *Tribolium* wing enhancer was active in *Drosophila*, we noticed that the activity of this enhancer was somewhat restricted, as it was active mainly at the dorsal-ventral (DV) compartmental boundary of the T2 wing, and only in a few cells in the haltere. This is in contrast with the expression in *Tribolium*, which showed a broader activity domain in the entire wing tissues both in the T2 and T3 segments. These differences in the activity domains suggest that some components that regulate the *Tribolium nub* wing enhancer are missing from the *Drosophila* T2 wing and almost entirely absent in the haltere. This highlights the limitation of inter-species analyses and the importance of performing reporter assays in the native species. The reporter assay system we developed now allows us to analyze enhancer activities in *Tribolium*. The successful demonstration of reporter analyses for *nub* in the wing and *hb* in the embryo suggest that our reporter construct works in various tissues; however, it is still crucial to evaluate the applicability of this system in diverse contexts.

### Choice of core promoters in reporter constructs

Our study showed that the choice of promoters is critical when assessing enhancer activity. *Tc-hsp68* was our first choice because it has successfully been used in the Gal4/UAS system in *Tribolium* (Schinko et al., 2010). However, in our reporter assay, although this promoter worked efficiently in *Drosophila*, it failed to drive reporter expression even with a functional enhancer in *Tribolium* (at least in our hands). Interestingly, the transgenic beetles with the *Tc-hsp68* reporter construct showed high occurrence of enhancer trap events (Fig. 4F-M), even though this promoter failed to work with the enhancer we placed right upstream of it. One explanation is that this promoter requires a certain distance for optimal interaction with enhancers in *Tribolium*. The situation might be less strict in *Drosophila* (for an unknown reason), allowing the *Tc-hsp68* promoter to overcome the distance requirement.

We also tried to assess the *nub* wing enhancer activity with the *nub* endogenous promoter, but to our surprise, this construct did not drive any expression. There are several possible explanations for this outcome. First, the region we selected might not contain the correct promoter for the *nub* transcript, although our 5’ RACE results (as well as the published *Tribolium* genome annotation (Tribolium Genome Sequencing et al., 2008)) supports our annotation of the *nub* transcription start site (Clark-Hachtel et al., 2013). Second, the 2kb region we used as the promoter may contain a suppressor element, interfering with the enhancer to drive reporter expression. Third, the *nub* promoter might require a long distance to interact properly with the wing enhancer, as the wing enhancer we identified is 40kb away from the *nub* transcription start site. This might parallel the situation with *Tc-hsp68*, in which this promoter preferably works with enhancers located at a certain distance. This may further support the idea that *Drosophila* are more permissive to changes in the enhancer/promoter distance. However, in the case of the *nub* endogenous promoter, there might be additional issues other than enhancer/promoter distance that prevented this reporter construct from working even in *Drosophila.*

The reporter construct with the DSCP (piggyGUM) worked efficiently both in *Drosophila* and *Tribolium*. The DCSP is a synthetic core promoter, composed of several common core promoter motifs (i.e. TATA box, Inr, MTE, and DPE) isolated from the *Drosophila* genome. The DSCP has been shown to work efficiently with a diverse array of developmental enhancers in various contexts in *Drosophila* (Pfeiffer et al., 2008; Zabidi et al., 2015), suggesting that this promoter may also work well with other enhancers in *Tribolium*. However, it is worth mentioning that a synthetic promoter similar to the DCSP, SCP1 (composed of *Drosophila* and viral promoter motifs (Juven-Gershon et al., 2006)), failed to work when tested in the Gal4/UAS system in *Tribolium* (Schinko et al., 2010). This again emphasizes the importance of choosing the correct promoter that fits the context of the study, which remains a critical area for further exploration.

### Enhancer studies in evo-devo

The study of enhancers and other CREs is critical to understand the molecular basis underlying morphological evolution, as changes in gene regulation, rather than the acquisition of new genes or the modification of protein structures, are often responsible for the evolution of the diverse array of morphology (Carroll, 2008). For example, changes in enhancers can facilitate evolution of novel structures via co-opting preexisting GRNs into a new context. Acquisition of enhancers *de novo* may also play a critical role in morphological novelty. Therefore, studying both evolutionarily conserved and diverged enhancers will help further our understanding of morphological evolution (see (Monteiro and Podlaha, 2009) for a comprehensive discussion of how *cis* studies can help elucidate the molecular basis for the evolution of novel traits). However, it has been a challenge to study enhancers in non-traditional model insects due to the lack of a reliable enhancer identification strategy. In this study, we showed that FAIRE-seq is readily applicable to non-traditional model species. Furthermore, the DSCP can be a useful promoter for establishing a reporter assay system and investigating the evolution of enhancers in non-*Drosophila* insects. Therefore, FAIRE-based chromatin profiling, along with reporter assay systems applicable to various insects, will make the research on enhancers more accessible, which will provide us with more insights into the evolution of the regulatory mechanisms underlying morphological diversity.

## MATERIALS AND METHODS

### Fly stocks

The following two *Drosophila* strains used in this study were obtained from the Bloomington *Drosophila* Stock center.

*P{UAS-Dcr-2.D}^1^, w^1118^; P{GawB} nubbin-AC-62*

*y*^*1*^ *w ^*^; wg ^Sp-1^/CyO, P{Wee-P.ph0}Bacc^Wee-P20^; P{20XUAS-6XGFP}attP2*.

### Beetle cultures

The beetle cultures were reared on whole wheat flour (+5% yeast) at 30 °C in a temperature and humidity controlled incubator. The *nub* enhancer trap line *pu11*, which has enhanced yellow fluorescent protein (EYFP) expression in the hindwing and elytron discs (Clark-Hachtel et al., 2013; Lorenzen et al., 2003; Tomoyasu et al., 2005), was used as to monitor *nub* expression in *Tribolium*.

### Tissue preparation for FAIRE

For the *Tribolium* larval T2 and T3 wing tissues, the dorso-lateral portion of the epidermal tissues that contain elytron (T2) and hindwing (T3) discs were dissected from the last instar larvae. Although these samples largely consisted of tissues that give rise to wing structures, they also contained body wall tissues as well as larval muscles due to difficulty of precisely dissecting the wing tissues from larvae. About 50 larvae (100 dissected tissues) were used for each biological replicate, with three replicates prepared for each wing sample. The brains were dissected from the head of the last instar larvae. About 40 brains were used for each biological replicate, with two replicates prepared. Embryos were collected in whole wheat flour (+5% yeast) for 24 hours at 30 °C. The collected embryos were cultured for one and two days at 30 °C for the 24-48 hour and 48-72 hour samples, respectively. 0.1g of embryos were used for each biological replicate, with three replicates prepared for each sample. These tissues and embryos were crosslinked with 4% formaldehyde for 30 min (larval tissues) or 8% formaldehyde for 30 min (embryos).

### FAIRE-seq analysis

FAIRE was performed as previously described (McKay and Lieb, 2013). FAIRE-seq libraries were sequenced on an Illumina HiSeq 2000 at the University of North Carolina High-Throughput Sequencing Facility. 50bp single-end Illumina reads were obtained for FAIRE-treated samples and two non-FAIRE-treated input samples. Reads were trimmed to remove index sequence and mapped to the *Tribolium* reference genome (version 3.0) with bowtie2 (Langmead and Salzberg, 2012). Read alignments were quality filtered (Q<10 dropped) and duplicate reads were removed using SAMtools. For visualization of FAIRE signal, bigwig files were produced by merging tissue/stage-specific replicate bam files with SAMtools and normalizing reads to sequencing depth using deepTools. These files were then visualized with the IGV genome viewer (Robinson et al., 2011; Thorvaldsdóttir et al., 2013). Peaks were called on individual replicates using MACS2 with the merged input sample bam files as the control. The *Drosophila* FAIRE profiles used in this study were previously published (McKay and Lieb, 2013). For differentially open peak analysis, mapped reads (.bam files) for each replicate and the merged input, along with MACS2 peaks (.narrowPeak files) called for each replicate, were provided as input for DiffBind. DiffBind creates a consensus peakset for all replicates provided, requiring a consensus peak to be present in at least 2 replicates of 1 sample. An experiment-wide consensus peakset was produced using all samples. Pairwise analysis of differentially open peaks between samples was performed within DiffBind with the DESeq2 method for all consensus peaksets, and plotted using the dba.plotMA() function. The differentially open peaks are listed in Table S6.

### Genome-wide GC-contents analysis

Using the experiment-wide consensus peakset described above, 1 kb of sequence upstream and downstream of each peak center was extracted from the genome using BEDTools (Quinlan and Hall, 2010) and custom Python scripts. For these 2kb fragments, those free of Ns were subjected to GC analysis. Changes in local GC content (250bp sliding window, 10bp step) were plotted against the whole-fragment average of GC content for all fragments. For the GC-rich region distance analysis, first, bedGraphs of GC content fluctuations above and below the genome wide average were computed at 70 and 60bp resolution for the *Tribolium* and *Drosophila* genomes, respectively. The genome of *Bombyx mori*, as well as the genomes of several coleopteran insects (*Agrilus planipennis*, *Dendroctonus ponderosae*, *Anoplophora glabripennis*, *Leptinotarsa decemlineata*, *Nicrophorus vespilloides*, and *Onthophagus taurus*) were analyzed at 70bp resolution. Peaks were then called using the bdgcallpeak command in MACS2. Distance between the edges of adjacent peaks was categorized into 100bp bins and the ln of the number of occurrences plotted. For the FAIRE peak distance analysis, distances between FARIE peaks were collected and plotted in the same manner as the GC peaks. A consensus *Drosophila* FAIRE peakset was obtained from DiffBind with the same settings as the *Tribolium* data using the previously published data (McKay and Lieb, 2013).

### Comparison between FAIRE and SCRMshaw

Enhancers predicted by SCRMshaw were taken from Table S4 of Kazemian et al. (Kazemian et al., 2014) and converted into BED format. BEDTools *merge* was used to combine overlapping and/or redundant (i.e., from more than one SCRMshaw scoring method) predictions, reducing the total number of predicted enhancers to 1214. BEDTools *intersect* was then used to determine all predicted enhancers with at least 50 bp overlap with a FAIRE peak (-f 0.10). FAIRE peaks not assigned to a *Tribolium* chromosome (i.e., not starting with “ChLG”) were omitted. Significance of overlaps was determined using BEDTools *fisher;* all overlaps were highly significant with –log(*P*) ≥ 19. Because this method provides only an approximation, a selection of datasets was tested via randomization. BEDTools *shuffle* was used to generate 1000 random intervals and the intersections were determined as above. The mean and standard deviation of the randomized intersections were calculated and used with the observed (SCRMshaw) intersection value to determine a *z* score. *P* values from all randomization tests were highly significant.

### *Drosophila* reporter assay constructs

pFUGG, a *Drosophila* GATEWAY-compatible phiC31 transformation plasmid, was used for reporter assay in *Drosophila* (McKay and Lieb, 2013). The phiC31 system allows site-specific integration (Bischof et al., 2007), thus preventing position effects due to different insertion sites. An enhancer cloned into pFUGG will drive Gal4 as the reporter, whose expression domains will then be visualized by crossing to UAS-EGFP flies.

### GATEWAY compatible piggyBac reporter constructs

The piggyBac plasmid with the 3xP3-EGFP marker construct and the FseI/AscI cloning site (Horn and Wimmer, 2000) was used to make all piggyBac constructs used in this study. For **piggyGHR** (<u>piggy</u>Bac <u>G</u>ATEWAY Tc-<u>h</u>sp68 ds<u>R</u>ed), the *gypsy* element, *Tc-hsp68* core promoter, *dsRed*, and the SV40 polyA signal were amplified by PCR, assembled through ligation, and inserted into the FseI/AscI site of the piggyBac plasmid. The assembled plasmid was then converted to a GATEWAY compatible plasmid by Gateway® Vector Conversion System (ThermoFisher Science). For **piggyGUM** (<u>piggy</u>Bac <u>G</u>ATEWAY <u>U</u>niversal promoter <u>m</u>Cherry), the reporter construct including the GATEWAY cassette was amplified from a *Drosophila* GATEWAY-compatible phiC31 transformation vector and inserted into the FseI/AscI site of the piggyBac plasmid. The primers used to make piggyGUM were listed in Table S5. The reporter constructs in **piggyNub-proR** (*piggy*Bac *nub* promoter dsRed) and **piggyAct5c-proR** (*piggy*Bac *Act5c* promoter dsRed) were *de novo* synthesized and inserted into the FseI/AscI site of the piggyBac plasmid.

### Enhancer cloning

Genomic fragments corresponding to possible enhancer regions were PCR amplified and cloned into pENTR using pENTR-D Directional TOPO Cloning kit (Thermo-Fisher Scientific, K240020). The primers used to clone the enhancer regions from the *Drosophila* and *Tribolium* genome are listed in Table S5. Cloned genomic fragments were then inserted into reporter constructs via GATEWAY Clonase reaction (Thermo-Fisher Scientific, 11791-019).

### *Drosophila* and *Tribolium* transgenesis

For *Drosophila* transgenesis, pFUGG constructs were transformed into the attP2 site (68A4) through PhiC31 integrase-mediated transgenesis system, and piggyBac constructs were transformed into *w*^*1118*^ with EGFP as a visible marker (BestGene *Drosophila* transgenic service). For *Tribolium* transgenesis, piggyBac constructs were transformed into *vermilion ^white^* with EGFP as a visible marker (TriGenES *Tribolium* Genome Editing Service for the *nub* and *Act5c* constructs, Friedrich-Alexander-Universität Erlangen-Nürnberg for the *hb* construct).

### Immunohistochemistry and *in situ* hybridization

*Drosophila* imaginal discs were dissected from the third instar larvae and fixed with 4% formaldehyde for 25 min. *Tribolium* elytron and hindwing discs were dissected from the last instar larvae, and fixed with 4% formaldehyde for 25 min. Dissected tissues were then washed and blocked with 10% BSA, and incubated with Rabbit anti-mCherry antibody (1:500; Abcam, ab167453) at 4 °C for overnight. After washing for one hour, the tissues were incubated with the Alexa 555 conjugated Goat anti-Rabbit antibody (1:500) for 2 hours at room temperature. All the discs were mounted on glass slides with ProLong® Gold antifade reagent (Life Technologies) for documentation. *in situ* hybridization was performed as previously described (Shippy et al., 2009), with DIG-labeled riboprobes and alkaline phosphatase conjugated anti-DIG antibody (Sigma-Aldrich 11093274910). Signal was developed using BM Purple (Sigma-Aldrich 11442074001). The primers used to amply the *mCherry* fragment for riboprobe synthesis were included in Table S5. The *hb* riboprobe used in this study is previously described (Wolff et al., 1998).

### Image Processing and Documentation

The images were captured by Zeiss 710 confocal microscope (mounted discs) and Zeiss AxioCam MRc5 with Zeiss Discovery V12 (*Tribolium* larvae and pupae). A filter set specific to mCherry (575/50x, 640/50m) was used to visualize the mCherry expression driven by piggyGUM constructs. *Tribolium* germband embryos were imaged with ProgRes CFcool camera on Zeiss Axio Scope.A1 microscope using ProgRes CapturePro image acquisition software. Some pictures were enhanced only for brightness and contrast with Adobe Photoshop.

## Acknowledgements

We thank the Bloomington Stock Center for fly stocks, and the Center for Bioinformatics and Functional Genomics (CBFG) and Center for Advanced Microscopy and Imaging (CAMI) at Miami University for technical support. We also thank Shuxia Yi for technical support, and Courtney Clark-Hachtel, David Linz, and other members of Tomoyasu lab for helpful discussion.

## Author contributions

Y.T. designed the project. Y-T.L., K.D.D., F.B-C., N.S., K.S., M.S.H., D.J.M., Y.T. performed experiments. Y-T.L., K.D.D., F.B-C., N.S., K.S., M.S.H., D.J.M., Y.T. analyzed the data. Y-T.L., K.D.D., M.S.H, D.J.M, and Y.T., wrote the manuscript.

## Data availability

FAIRE-seq data have been deposited at Gene Expression Omnibus (GEO) under accession number GSE104495.

## SUPPLEMENTAL FIGURES

**Fig. S1.**
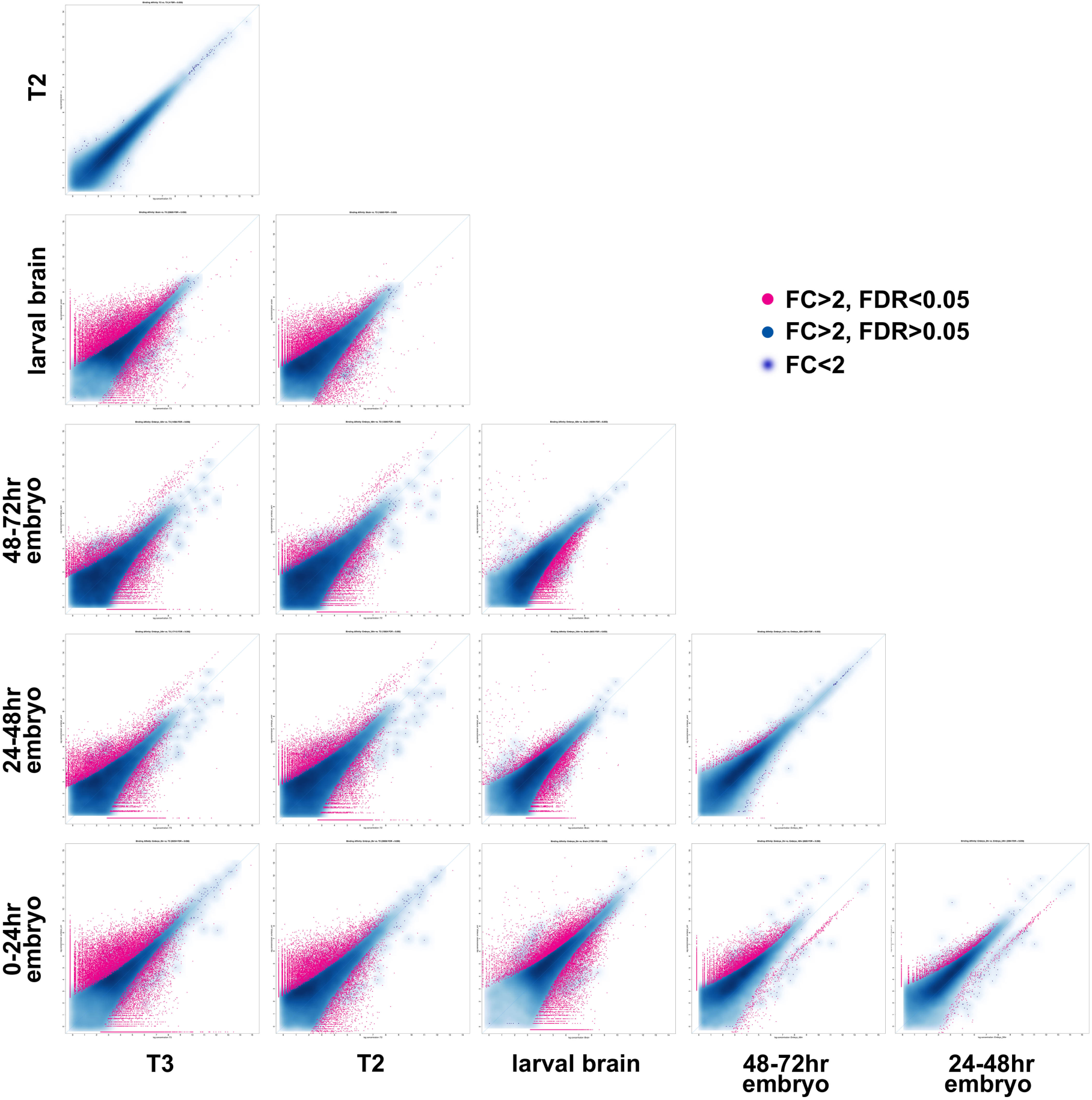
Differentially open peak analysis. Pairwise analysis of differentially open peaks between samples. Red represents peaks that exhibit over two-fold change (FC) between samples with the false discovery rate (FDR) < 0.05, while blue represents peaks over two-fold change but with FDR > 0.05. The blue cloud represents peaks with less than two-fold change between samples.

**Fig. S2.**
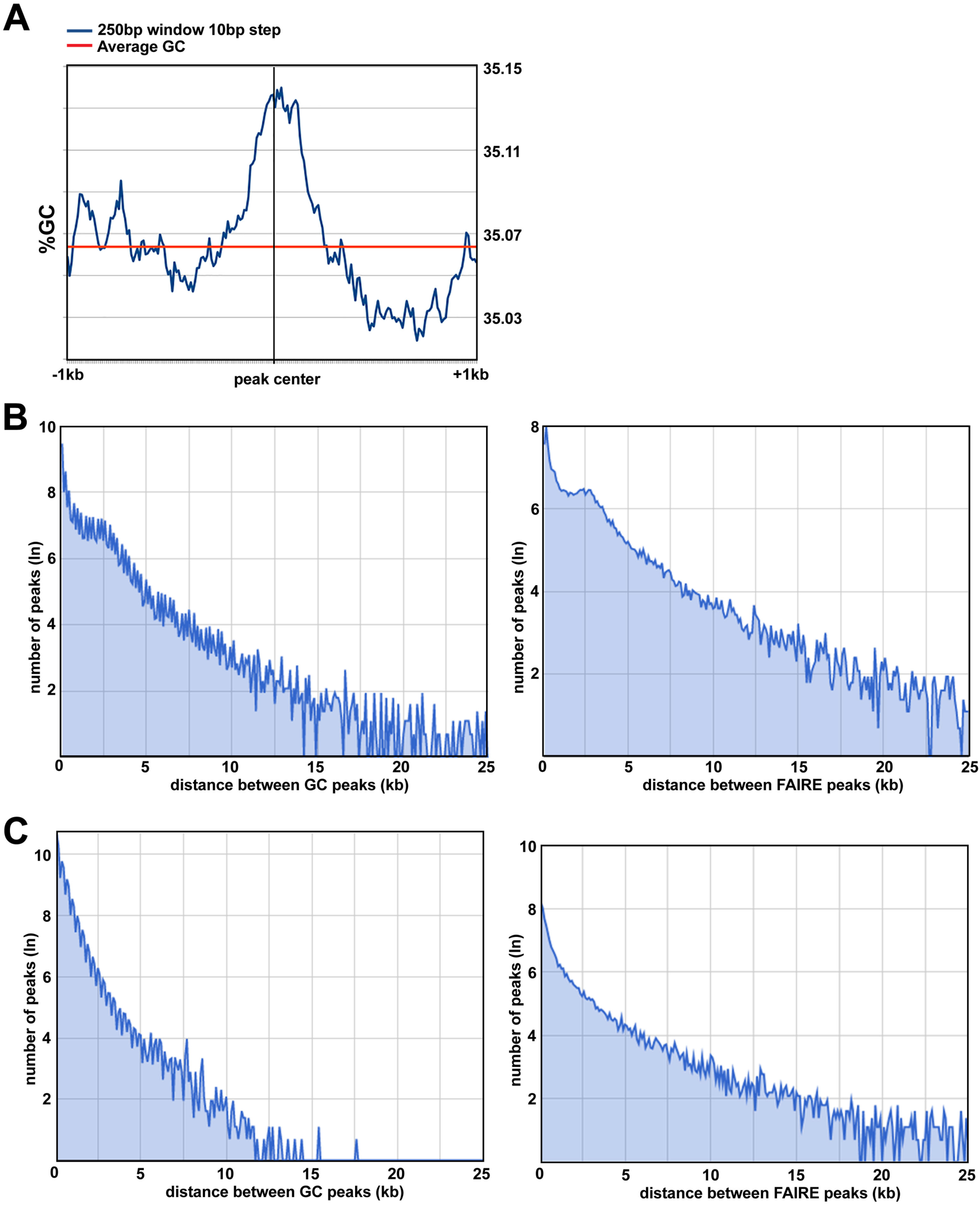
Distribution of FAIRE peaks and GC-rich regions. (A) Correlation between FAIRE peaks and high GC contents in *Tribolium*. (B) Distribution of intervals between FAIRE peaks in *Tribolium* and *Drosophila*. (C) Distribution of intervals between GC-rich regions in *Tribolium* and *Drosophila*. Note that there is a significant accumulation around 3 kb in *Tribolium* but not in *Drosophila* (B, C).

**Fig. S3.**
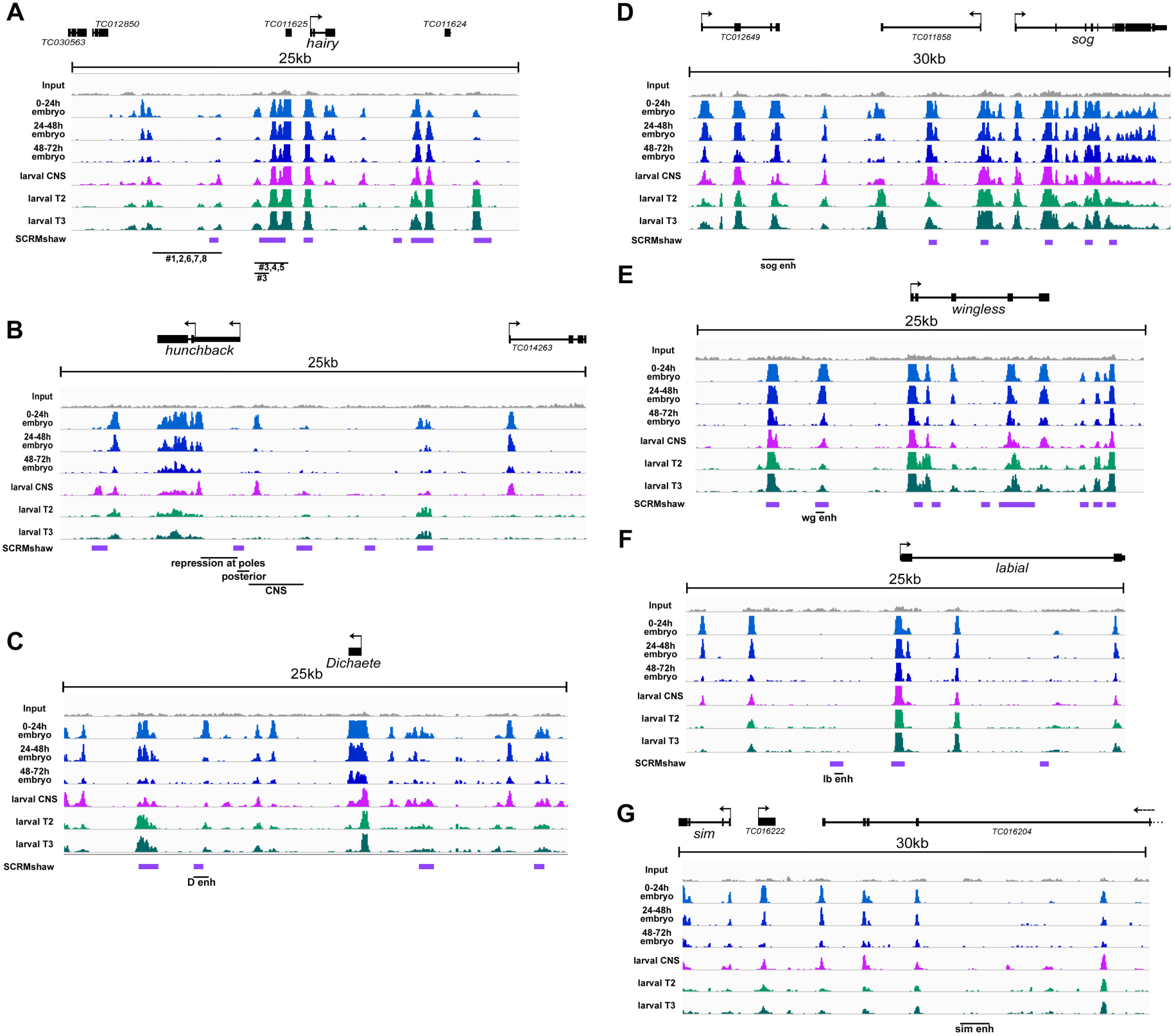
Comparison of the FAIRE data to previous enhancer studies. (A) *hairy* (*h*), (B) *hunchback* (*hb*), (C) *Dichaete* (*D*), (D) *short gastrulation* (*sog*), (E) *wingless* (*wg*), (F) *labial* (*lab*), (G) *single-minded* (*sim*). Previously described possible enhancer regions at these loci are shown by black lines underneath the FAIRE profiles. SCRMshaw predictions are also shown (purple). Only the enhancers at the *h* locus have been tested in the native *Tribolium* context, while other enhancers were evaluated in the cross-species context. FAIRE peaks match well with the previously described enhances for *h*, *hb*, *D*, *sog*, and *wg* (A-E), while no significant overlaps are observed between FAIRE peaks and the previously described enhancers for *lab* and *sim*.

**Fig. S4.**
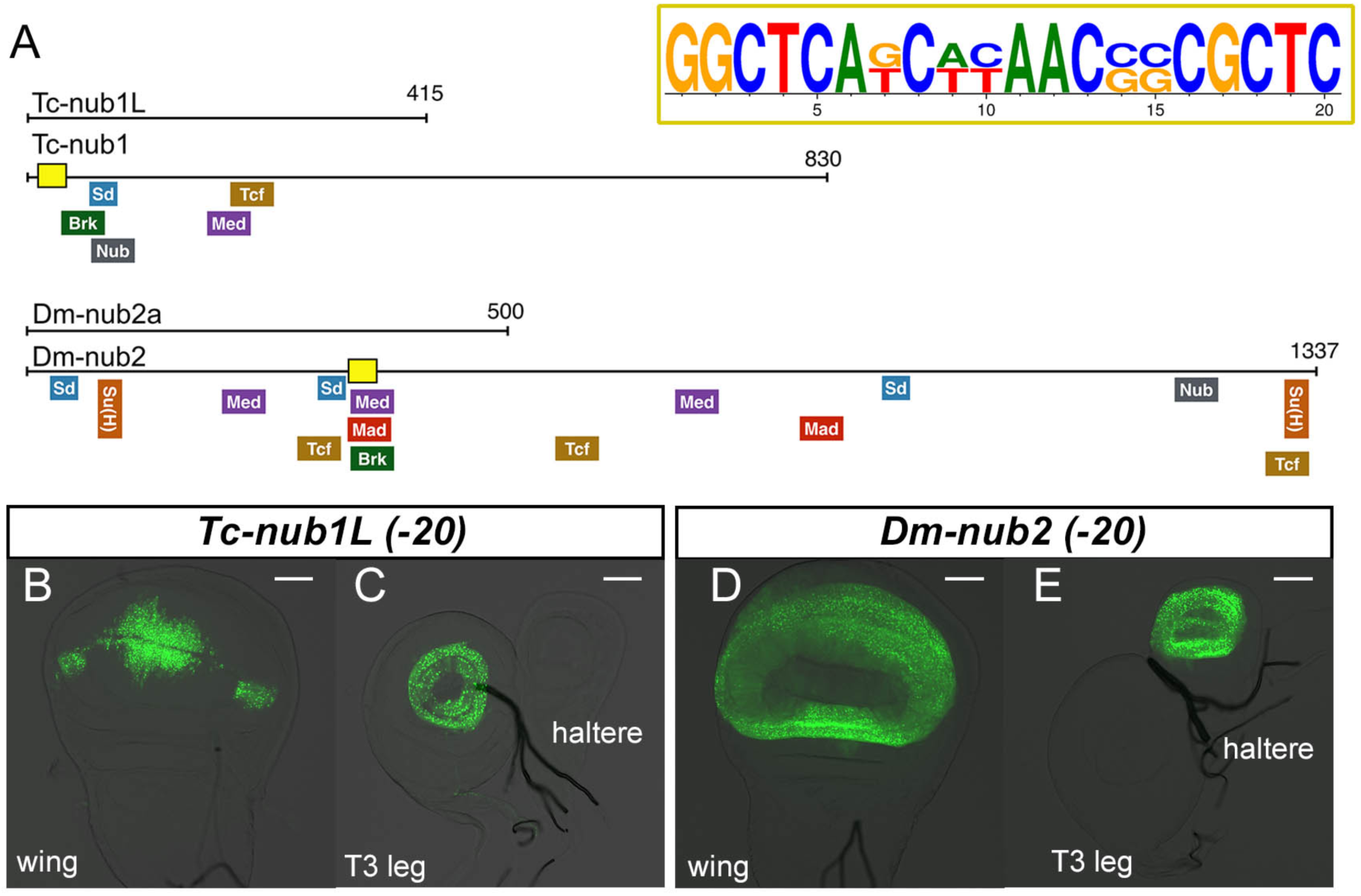
Deletion of the motif shared between the *Tribolium* and *Drosophila nub* wing enhancers. (A) Locations of TF binding sites within the *Tribolium* and *Drosophila nub* wing enhancers. A 20bp shared sequence between the two enhancers is shown with a yellow box. (B, C) Activities of the *Tribolium* and *Drosophila nub* wing enhancers when the conserved 20bp shared sequence is deleted. No significant changes in enhancer activity are observed, indicating that the 20bp sequence is dispensable from the activity of the two enhancers. Scale bar: 50 µm.

## SUPPLEMENTAL TABLES

**Table S1.** Overlap between SCRMshaw predictions and FAIRE peaks

| Training_set <sup>a</sup> | n <sup>b</sup> | FAIRE<br>0-24hr | FAIRE<br>24-48hr | FAIRE<br>48-72hr | FAIRE<br>larvalT2 | FAIRE<br>larvalT3 | combined<br>(embryo) | combined<br>(larval) | combined |
| --- | --- | --- | --- | --- | --- | --- | --- | --- | --- |
| ap | 129 | 60.5% | 55.8% | 55.0% | 73.6% | 72.1% | nd<sup>c</sup> | nd | nd |
| blastoderm | 138 | 76.1% | 72.5% | 68.8% | 87.0% | 89.9% | nd | nd | nd |
| cns | 256 | 86.7% | 82.0% | 83.6% | 93.4% | 93.0% | nd | nd | nd |
| dorsal_ectoderm | 183 | 79.2% | 74.9% | 74.3% | 88.0% | 87.4% | nd | nd | nd |
| dv | 137 | 69.3% | 65.0% | 67.2% | 86.9% | 84.7% | nd | nd | nd |
| ectoderm+ectoderm.2 | 280 | 76.4% | 73.6% | 74.6% | 86.8% | 88.2% | nd | nd | nd |
| endoderm | 109 | 68.8% | 63.3% | 66.1% | 78.0% | 82.6% | nd | nd | nd |
| imaginal_disc (1+2) | 246 | 85.4% | 83.3% | 81.3% | 93.9% | 94.3% | nd | nd | nd |
| mesectoderm | 42 | 83.3% | 81.0% | 81.0% | 97.6% | 92.9% | nd | nd | nd |
| somatic_muscle | 42 | 78.6% | 76.2% | 81.0% | 83.3% | 88.1% | nd | nd | nd |
| ventral_ectoderm | 129 | 89.1% | 86.8% | 86.8% | 90.7% | 92.2% | nd | nd | nd |
| wing.2 | 99 | 91.9% | 90.9% | 90.9% | 98.0% | 98.0% | nd | nd | nd |
| **all combined** | **1214** | **74.1%** | **70.5%** | **71.1%** | **85.1%** | **85.3%** | **78.8%** | **88.1%** | **90.3%** |

<sup>a</sup> Training sets as listed in Kazemian et al. (2014). "all\_combined" is all 1214 enhancer predictions. Not all individual training sets are shown.
<sup>b</sup> Total predicted enhancers
<sup>c</sup> nd: not done

**Table S2.** The list of FARIE peaks that overlap SCRMshaw. (attached separately)

| Enhancer | wing | haltere | leg | antennae<br>/eyes | CNS* | trachea | gut | mouthpart | salivary<br>gland* | others |
| --- | --- | --- | --- | --- | --- | --- | --- | --- | --- | --- |
| *Tc-nub1* | DV<br>boundary | a few cells | + | antennae &<br>optical nerves | ++ | + | ++ | + | - | a circle of cells near the<br>posterior end |
| *Tc-nub2* | - | - | + | antennae | + | - | + | + | + |  |
| *Tc-nub3* | notum | notum | - | - | + | - | ++ | + | + | spiracles |
| *Tc-nub1Core* | - | - | - | - | + | - | + | + | + | a line of cells in epidermis |
| *Tc-nub1L* | DV<br>boundary | a few cells | + | antennae &<br>optical nerves | ++ | + | + | + | - |  |
| *Tc-nub1R* | - | - | - | - | + | - | ++ | - | + | spiracles, a circle of cells<br>near the posterior end |
| *Tc-nub1a* | DV<br>boundary | a few cells | + | antennae &<br>optical nerves | ++ | - | ++ | + | + |  |
| *Tc-nub1Lb* | DV<br>boundary | a few cells | + | antennae &<br>optical nerves | ++ | + | ++ | + | + |  |

\*CNS and salivary gland expression may not be specific to the construct tested.

**Table S3.** Expression of *Tc nub* reporter constructs outside of the wing and leg imaginal disc in *Drosophila*

| Enhancer | wing | haltere | leg | antennae<br>/eyes | CNS | trachea | gut | mouthpart | Salivary<br>gland |
| --- | --- | --- | --- | --- | --- | --- | --- | --- | --- |
| *Dm-nub1* | - | - | - | - | ++ | - | ++ | - | + |
| *Dm-nub2* | pouch | pouch | - | - | ++ | - | ++ | + | + |
| *Dm-nub3* | - | - | ++ | - | ++ | - | ++ | + | + |
| *Dm-nub2a* | pouch | pouch | - | + | + | - | + | + | - |
| *Dm-nub2b* | - | - | - | - | + | - | + | - | - |
| *Dm-nub2c* | - | - | - | - | + | - | ++ | - | - |

\*CNS and salivary gland expression may not be specific to the construct tested.

**Table S4.**
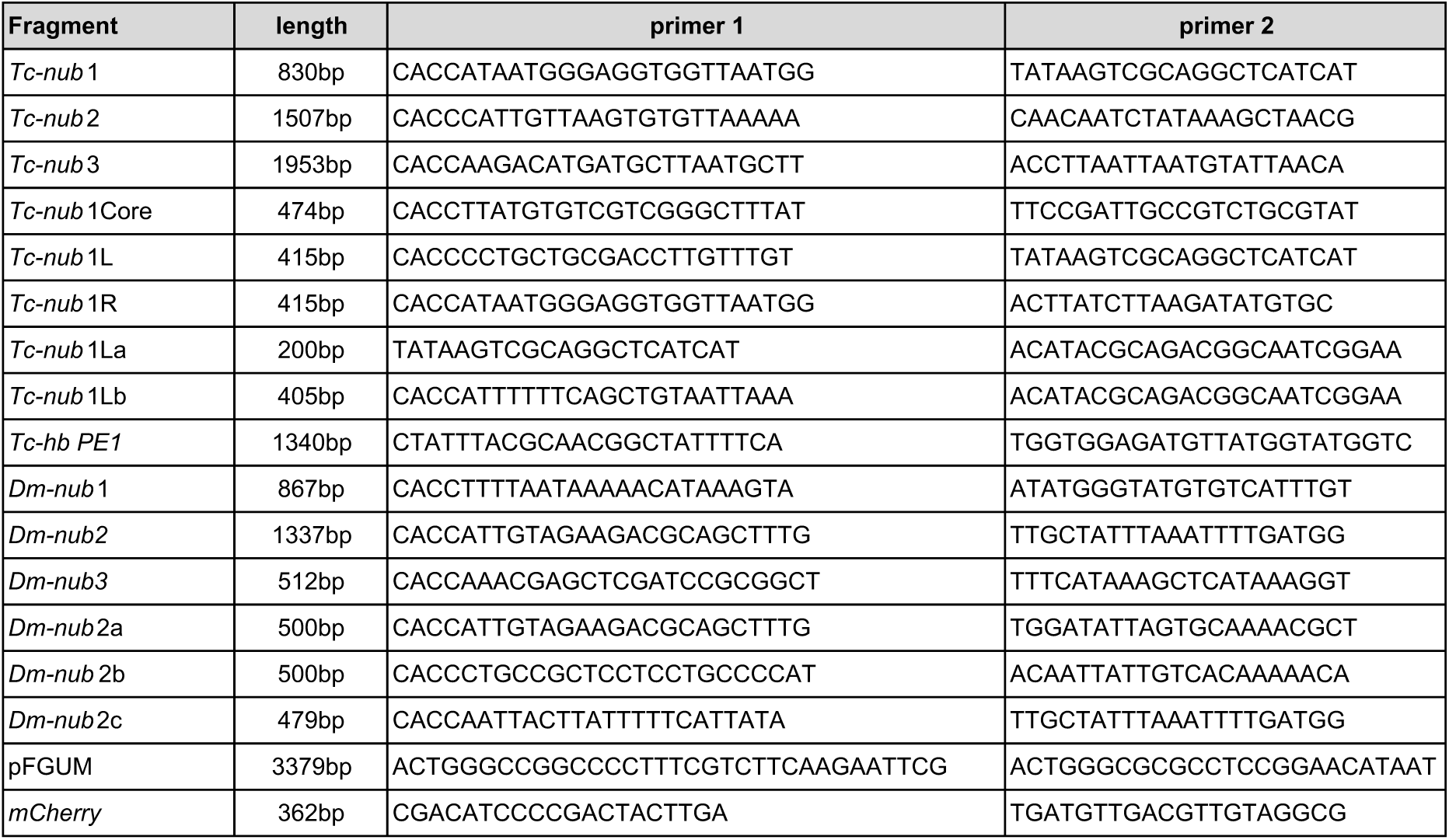
Expression of *Dm nub* reporter constructs outside of the wing and leg imaginal disc in *Drosophila*

**Table S5. Primers used in this study**

**Table S6. The list of differentially accessible FARIE peaks.** (attached separately)

## REFERENCES

Angelini, D. R., Smith, F. W. and Jockusch, E. L. (2012). Extent With Modification: Leg Patterning in the Beetle Tribolium castaneum and the Evolution of Serial Homologs. G3 2, 235–248.

Belles, X. (2010). Beyond Drosophila: RNAi in vivo and functional genomics in insects. Annu Rev Entomol 55, 111–128.

Bischof, J., Maeda, R. K., Hediger, M., Karch, F. and Basler, K. (2007). An optimized transgenesis system for Drosophila using germ-line-specific phiC31 integrases. Proc. Natl. Acad. Sci. U. S. A. 104, 3312–7.

Brown, S. J., Mahaffey, J. P., Lorenzen, M. D., Denell, R. E. and Mahaffey, J. W. (1999). Using RNAi to investigate orthologous homeotic gene function during development of distantly related insects. Evol Dev 1, 11–15.

Bucher, G., Scholten, J. and Klingler, M. (2002). Parental RNAi in Tribolium (Coleoptera). Curr Biol 12, R85–6.

Cande, J., Goltsev, Y. and Levine, M. S. (2009). Conservation of enhancer location in divergent insects. Proc. Natl. Acad. Sci. 106, 14414–14419.

Carroll, S. B. (2008). Evo-Devo and an Expanding Evolutionary Synthesis: A Genetic Theory of Morphological Evolution. Cell 134, 25–36.

Carroll, S., Grenier, J. K. and Weatherbee, S. D. (2005). From DNA to Diversity. second edi. Blackwell publishing.

Chung, Y. T. and Keller, E. B. (1990). Positive and negative regulatory elements mediating transcription from the Drosophila melanogaster actin 5C distal promoter. Mol. Cell. Biol. 10, 6172–80.

Clark-Hachtel, C. M., Linz, D. M. and Tomoyasu, Y. (2013). Insights into insect wing origin provided by functional analysis of vestigial in the red flour beetle, Tribolium castaneum. Proc Natl Acad Sci U S A 110, 16951–16956.

Denell, R. (2008). Establishment of tribolium as a genetic model system and its early contributions to evo-devo. Genetics 180, 1779–1786.

Eckert, C., Aranda, M., Wolff, C. and Tautz, D. (2004). Separable stripe enhancer elements for the pair-rule gene hairy in the beetle Tribolium. EMBO Rep. 5, 638–42.

Frazer, K. A., Pachter, L., Poliakov, A., Rubin, E. M. and Dubchak, I. (2004). VISTA: computational tools for comparative genomics. Nucleic Acids Res. 32, W273–W279.

Giresi, P. G., Kim, J., McDaniell, R. M., Iyer, V. R. and Lieb, J. D. (2007). FAIRE (Formaldehyde-Assisted Isolation of Regulatory Elements) isolates active regulatory elements from human chromatin. Genome Res. 17, 877–85.

Halfon, M. S. (2017). Perspectives on Gene Regulatory Network Evolution. Trends Genet.

Hong, J.-W., Hendrix, D. A. and Levine, M. S. (2008). Shadow enhancers as a source of evolutionary novelty. Science 321, 1314.

Horn, C. and Wimmer, E. A. (2000). A versatile vector set for animal transgenesis. Dev. Genes Evol. 210, 630–7.

Jenett, A., Rubin, G. M., Ngo, T.-T. B., Shepherd, D., Murphy, C., Dionne, H., Pfeiffer, B. D., Cavallaro, A., Hall, D., Jeter, J., et al. (2012). A GAL4-driver line resource for Drosophila neurobiology. Cell Rep. 2, 991–1001.

Jory, A., Estella, C., Giorgianni, M. W., Slattery, M., Laverty, T. R., Rubin, G. M. and Mann, R. S. (2012). A survey of 6,300 genomic fragments for cis-regulatory activity in the imaginal discs of Drosophila melanogaster. Cell Rep. 2, 1014–24.

Juven-Gershon, T., Cheng, S. and Kadonaga, J. T. (2006). Rational design of a super core promoter that enhances gene expression. Nat. Methods 3, 917–22.

Kantorovitz, M. R., Kazemian, M., Kinston, S., Miranda-Saavedra, D., Zhu, Q., Robinson, G. E., Göttgens, B., Halfon, M. S. and Sinha, S. (2009). Motif-blind, genome-wide discovery of cis-regulatory modules in Drosophila and mouse. Dev. Cell 17, 568–79.

Katzen, F. (2007). Gateway ® recombinational cloning: a biological operating system. Expert Opin. 2 Drug Discov. 2, 571–589.

Kazemian, M., Zhu, Q., Halfon, M. S. and Sinha, S. (2011). Improved accuracy of supervised CRM discovery with interpolated Markov models and cross-species comparison. Nucleic Acids Res. 39, 9463–72.

Kazemian, M., Suryamohan, K., Chen, J.-Y., Zhang, Y., Samee, M. A. H., Halfon, M. S. and Sinha, S. (2014). Evidence for deep regulatory similarities in early developmental programs across highly diverged insects. Genome Biol. Evol. 6, 2301–20.

Kvon, E. Z., Kazmar, T., Stampfel, G., Yáñez-Cuna, J. O., Pagani, M., Schernhuber, K., Dickson, B. J. and Stark, A. (2014). Genome-scale functional characterization of Drosophila developmental enhancers in vivo. Nature 512, 91–5.

Li, L., Zhu, Q., He, X., Sinha, S. and Halfon, M. S. (2007). Large-scale analysis of transcriptional cis-regulatory modules reveals both common features and distinct subclasses. Genome Biol 8, R101.

Lorenzen, M. D., Berghammer, A. J., Brown, S. J., Denell, R. E., Klingler, M. and Beeman, R. W. (2003). piggyBac-mediated germline transformation in the beetle Tribolium castaneum. Insect Mol Biol 12, 433–440.

Lynch, J. A., El-Sherif, E. and Brown, S. J. (2012). Comparisons of the embryonic development of Drosophila, Nasonia, and Tribolium. Wiley Interdiscip. Rev. Dev. Biol. 1, 16–39.

Marques-Souza, H. (2007). Evolution of the gene regulatory network controlling trunk segmentation in insects.

Marques-Souza, H., Aranda, M. and Tautz, D. (2008). Delimiting the conserved features of hunchback function for the trunk organization of insects. Development 135, 881–888.

Mayor, C., Brudno, M., Schwartz, J. R., Poliakov, A., Rubin, E. M., Frazer, K. A., Pachter, L. S. and Dubchak, I. (2000). VISTA?: visualizing global DNA sequence alignments of arbitrary length. Bioinformatics 16, 1046–7.

McKay, D. J. and Lieb, J. D. (2013). A common set of DNA regulatory elements shapes Drosophila appendages. Dev. Cell 27, 306–18.

Monteiro, A. and Podlaha, O. (2009). Wings, Horns, and Butterfly Eyespots: How Do Complex Traits Evolve? PLoS Biol. 7, e1000037.

Ng, M., Diaz-Benjumea, F. J. and Cohen, S. M. (1995). Nubbin encodes a POU-domain protein required for proximal-distal patterning in the Drosophila wing. Development 121, 589–599.

Pearson, J. C., McKay, D. J., Lieb, J. D. and Crews, S. T. (2016). Chromatin profiling of Drosophila CNS subpopulations identifies active transcriptional enhancers. Development 143, 3723–3732.

Peel, A. D. (2008). The evolution of developmental gene networks: lessons from comparative studies on holometabolous insects. Philos. Trans. R. Soc. B Biol. Sci. 363, 1539–1547.

Pfeiffer, B. D., Jenett, A., Hammonds, A. S., Ngo, T.-T. B., Misra, S., Murphy, C., Scully, A., Carlson, J. W., Wan, K. H., Laverty, T. R., et al. (2008). Tools for neuroanatomy and neurogenetics in Drosophila. Proc. Natl. Acad. Sci. U. S. A. 105, 9715–20.

Quinlan, A. R. and Hall, I. M. (2010). BEDTools: a flexible suite of utilities for comparing genomic features. Bioinformatics 26, 841–2.

Robinson, J. T., Thorvaldsdóttir, H., Winckler, W., Guttman, M., Lander, E. S., Getz, G. and Mesirov, J. P. (2011). Integrative genomics viewer. Nat. Biotechnol. 29, 24–26.

Schinko, J. B., Weber, M., Viktorinova, I., Kiupakis, A., Averof, M., Klingler, M., Wimmer, E. A. and Bucher, G. (2010). Functionality of the GAL4/UAS system in Tribolium requires the use of endogenous core promoters. BMC Dev Biol 10, 53.

Schmitt-Engel, C., Schultheis, D., Schwirz, J., Ströhlein, N., Troelenberg, N., Majumdar, U., Dao, V. A., Grossmann, D., Richter, T., Tech, M., et al. (2015). The iBeetle large-scale RNAi screen reveals gene functions for insect development and physiology. Nat. Commun. 6, 7822.

Shippy, T. D., Coleman, C. M., Tomoyasu, Y. and Brown, S. J. (2009). Concurrent in situ hybridization and antibody staining in red flour beetle (Tribolium) embryos. Cold Spring Harb Protoc 2009, pdb prot5257.

Shlyueva, D., Stampfel, G. and Stark, A. (2014). Transcriptional enhancers: from properties to genome-wide predictions. Nat. Rev. Genet. 15, 272–286.

Simon, J. M., Giresi, P. G., Davis, I. J. and Lieb, J. D. (2012). Using formaldehyde-assisted isolation of regulatory elements (FAIRE) to isolate active regulatory DNA. Nat. Protoc. 7, 256–267.

Smith, F. W., Angelini, D. R., Gaudio, M. S. and Jockusch, E. L. (2014). Metamorphic labral axis patterning in the beetle Tribolium castaneum requires multiple upstream, but few downstream, genes in the appendage patterning network. Evol. Dev. 16, 78–91.

Song, L., Zhang, Z., Grasfeder, L. L., Boyle, A. P., Giresi, P. G., Lee, B.-K., Sheffield, N. C., Gräf, S., Huss, M., Keefe, D., et al. (2011). Open chromatin defined by DNaseI and FAIRE identifies regulatory elements that shape cell-type identity. Genome Res. 21, 1757–67.

Sosinsky, A., Honig, B., Mann, R. S. and Califano, A. (2007). Discovering transcriptional regulatory regions in Drosophila by a nonalignment method for phylogenetic footprinting. Proc Natl Acad Sci U S A 104, 6305–6310.

Stark, A., Lin, M. F., Kheradpour, P., Pedersen, J. S., Parts, L., Carlson, J. W., Crosby, M. A., Rasmussen, M. D., Roy, S., Deoras, A. N., et al. (2007). Discovery of functional elements in 12 Drosophila genomes using evolutionary signatures. Nature 450, 219–232.

Suryamohan, K. and Halfon, M. S. (2015). Identifying transcriptional cis-regulatory modules in animal genomes. Wiley Interdiscip. Rev. Dev. Biol. 4, 59–84.

Thorvaldsdóttir, H., Robinson, J. T. and Mesirov, J. P. (2013). Integrative Genomics Viewer (IGV): high-performance genomics data visualization and exploration. Brief. Bioinform. 14, 178–92.

Tomoyasu, Y. (2017). Ultrabithorax and the evolution of insect forewing/hindwing differentiation. Curr. Opin. Insect Sci. 19,.

Tomoyasu, Y. and Denell, R. E. (2004). Larval RNAi in Tribolium (Coleoptera) for analyzing adult development. Dev Genes Evol 214, 575–578.

Tomoyasu, Y., Wheeler, S. R. and Denell, R. E. (2005). Ultrabithorax is required for membranous wing identity in the beetle Tribolium castaneum. Nature 433, 643–647.

Tomoyasu, Y., Arakane, Y., Kramer, K. J. and Denell, R. E. (2009). Repeated Co-options of Exoskeleton Formation during Wing-to-Elytron Evolution in Beetles. Curr. Biol. 19, 2057–2065.

Tribolium Genome Sequencing consortium, C., Richards, S., Gibbs, R. A., Weinstock, G. M., Brown, S. J., Denell, R., Beeman, R. W., Gibbs, R., Beeman, R. W., Brown, S. J., et al. (2008). The genome of the model beetle and pest Tribolium castaneum. Nature 452, 949–955.

Uyehara, C. M., Nystrom, S. L., Niederhuber, M. J., Leatham-Jensen, M., Ma, Y., Buttitta, L. A. and McKay, D. J. (2017). Hormone-dependent control of developmental timing through regulation of chromatin accessibility. Genes Dev. 31, 862–875.

Wang, L., Wang, S., Li, Y., Paradesi, M. S. and Brown, S. J. (2007). BeetleBase: the model organism database for Tribolium castaneum. Nucleic Acids Res 35, D476–9.

Wolff, C., Schröder, R., Schulz, C., Tautz, D. and Klingler, M. (1998). Regulation of the Tribolium homologues of caudal and hunchback in Drosophila: evidence for maternal gradient systems in a short germ embryo. Development 125, 3645–54.

Zabidi, M. A., Arnold, C. D., Schernhuber, K., Pagani, M., Rath, M., Frank, O. and Stark, A. (2015). Enhancer-core-promoter specificity separates developmental and housekeeping gene regulation. Nature 518, 556–9.

Zdobnov, E. M. and Bork, P. (2007). Quantification of insect genome divergence. Trends Genet. 23, 16–20.

Zhu, X., Rudolf, H., Healey, L., François, P., Brown, S. J., Klingler, M. and El-Sherif, E. (2017). Speed regulation of genetic cascades allows for evolvability in the body plan specification of insects. Proc. Natl. Acad. Sci. U. S. A. 201702478.

Zinzen, R. P., Cande, J., Ronshaugen, M., Papatsenko, D. and Levine, M. (2006). Evolution of the Ventral Midline in Insect Embryos. Dev. Cell 11, 895–902.

